# Scaled, high fidelity electrophysiological, morphological, and transcriptomic cell characterization

**DOI:** 10.1101/2020.11.04.369082

**Authors:** Brian R. Lee, Agata Budzillo, Kristen Hadley, Jeremy A. Miller, Tim Jarsky, Katherine Baker, DiJon Hill, Lisa Kim, Rusty Mann, Lindsay Ng, Aaron Oldre, Ram Rajanbabu, Jessica Trinh, Thomas Braun, Rachel Dalley, Nathan W. Gouwens, Brian E. Kalmbach, Tae Kyung Kim, Kimberly Smith, Gilberto J. Soler-Llavina, Staci A. Sorensen, Bosiljka Tasic, Jonathan T. Ting, Ed S. Lein, Hongkui Zeng, Gabe J. Murphy, Jim Berg

**Author notes:** These authors contributed equally (Brian R. Lee, Agata Budzillo). Correspondence (B.R.L.) and (A.B.).

## Abstract

The Patch-seq approach is a powerful variation of the standard patch clamp technique that allows for the combined electrophysiological, morphological, and transcriptomic characterization of individual neurons. To generate Patch-seq datasets at a scale and quality that can be integrated with high-throughput dissociated cell transcriptomic data, we have optimized the technique by identifying and refining key factors that contribute to the efficient collection of high-quality data. To rapidly generate high-quality electrophysiology data, we developed patch clamp electrophysiology software with analysis functions specifically designed to automate acquisition with online quality control. We recognized a substantial improvement in transcriptomic data quality when the nucleus was extracted following the recording. For morphology success, the importance of maximizing the neuron’s membrane integrity during the extraction of the nucleus was much more critical to success than varying the duration of the electrophysiology recording. We compiled the lab protocol with the analysis and acquisition software at https://github.com/AllenInstitute/patchseqtools. This resource can be used by individual labs to generate Patch-seq data across diverse mammalian species and that is compatible with recent large-scale publicly available Allen Institute Patch-seq datasets.

## Introduction

Describing and understanding the properties of neuronal cell types is a critical first step towards understanding circuit activity within the brain, and ultimately cognitive function. Neurons exhibit stereotyped yet diverse electrophysiological, morphological, and transcriptomic properties (Tasic et al. 2018; Zeng and Sanes 2017; Arendt et al. 2016; Kepecs and Fishell 2014; Tremblay, Lee, and Rudy 2016; Gouwens et al. 2019; Tasic et al. 2016; Gouwens et al. 2020) and understanding how each of these distinct features relate to one another may provide us with mechanistic insight into the roles of these neuron types. The introduction of single-cell RNA-sequencing (scRNA-seq) has revolutionized the field of transcriptomics (Zeisel et al. 2015). Dissociated cells or nuclei are isolated in a high-throughput manner to provide a comprehensive analysis of the molecular underpinnings of a single cell. Systematic and large-scale scRNA-seq approaches have been successful at characterizing brain cell types across mammalian species (Yao et al. 2020; Tasic et al. 2016; Tasic et al. 2018; Bakken et al. 2018; Hodge et al. 2019; Hashikawa et al. 2020; Zeisel et al. 2015; Mickelsen et al. 2019). These large-scale studies often include data from tens of thousands to millions of neurons whereas electrophysiological or morphological studies are limited to tens or hundreds of neurons. Despite scRNA-seq providing an in-depth look into gene expression and cell type classification, relationships to the morpho-electric neuron types described in the literature (Tremblay, Lee, and Rudy 2016) can only be inferred. Studies with triple modality data are rare and lack the scale to capture the true biological variability.

The Patch-seq recording technique is a powerful approach (Cadwell et al. 2015; Fuzik et al. 2015; Cadwell et al. 2017; van den Hurk et al. 2018) that can provide morphology (M), electrophysiology (E), and transcriptome (T) data from single cells, i.e. triple modality MET data. These data are an important tool towards establishing ‘correspondence’ of these properties through analysis. This technique is a modification of the well-established slice patch clamp electrophysiology approach, where intrinsic properties are recorded from neurons in acute brain slices while simultaneously filling the cell with biocytin. Following fixation, the biocytin is reacted with DAB as chromogen to generate a dark precipitate that fills the cell, enabling imaging and a digital morphological reconstruction. In the Patch-seq technique, the neuron’s cytoplasm is collected at the end of the recording, then processed via RNA-seq to identify its gene expression patterns. The technique has been used successfully to characterize both cortical and subcortical neurons (Berg 2020; Scala et al. 2020; Hashikawa et al. 2020; Gouwens et al. 2020; Muñoz-Manchado et al. 2018).

Despite its promise, early Patch-seq data suffered from three primary issues: 1) inconsistent quality, with non-specific cellular contamination and low yield of genes critical to accurate mapping to transcriptomic types (Tripathy et al. 2018), 2) little to no recovery of cell morphology (Cadwell et al. 2015), and 3) low throughput, with adequate cell fills requiring up to an hour of recording per cell. To address these issues we refined the Patch-seq technique to increase the efficiency of each step to minimize data attrition and increase the throughput to facilitate a more comprehensive analysis of cell type characterization (Berg 2020; Gouwens et al. 2020). We systematically modified the existing Patch-seq protocols, using feedback from experimental metadata to reveal the key determinants of success, including nucleus extraction and slow withdrawal of the recording electrode. We also used a novel electrophysiology acquisition platform, specialized to provide online quality control and adaptive stimuli designed to reduce experiment duration. Together with these adaptations, we demonstrate how high-quality triple modality information may be gathered across diverse brain cell types, different species and at scale.

Adopting the Patch-seq technique in an existing or new laboratory can be daunting due to the complexity of multiple data streams. To make adoption as simple as possible, we have created a resource, https://github.com/AllenInstitute/patchseqtools, as a starting point for labs interested in using the Patch-seq technique or refining their existing technique.

Specifically, this resource consists of three components: 1) a step-by-step optimized Patch-seq protocol, including helpful tips and a troubleshooting guide; 2) the Multichannel Igor Electrophysiology Suite (MIES) software package, including built-in analysis functions to increase the efficiency and robustness of Patch-seq data; and 3) an R library that uses a modified workflow from the patchSeqQC R library (Tripathy et al. 2018) to process and assay the quality of Patch-seq transcriptomic data. Here we highlight the components of this resource and detail the background behind critical protocol decisions. The resource balances detailed internal standards with flexibility to adjust to a specific user’s experimental approach. In the end, data generated using this resource can be benchmarked against the data from the over 7,000 publicly available Patch-seq neuron experiments that can be downloaded from https://portal.brain-map.org.

## Results

### A Patch-seq protocol optimized for fast, high quality data generation

We used broad and specific transgenic Cre driver mouse lines to target over 7,000 excitatory and inhibitory cells in adult mouse primary visual cortex (VISp). To compare neurons within similar neuron types, we focus our analysis on four established Cre lines: retinol binding protein 4 (Rbp4-Cre) for glutamatergic (excitatory) neurons and parvalbumin (Pvalb-Cre), somatostatin (Sst-Cre), and vasoactive intestinal peptide (Vip-Cre) for GABAergic (inhibitory) neurons. Each of these lines have been shown to be cell type- and/or brain region-specific (Madisen et al. 2009; Harris et al. 2014), and they label cells within the same transcriptomic class (e.g., glutamatergic, GABAergic) or subclass (e.g., Pvalb, Sst, Vip) (Tasic et al. 2016; Tasic et al. 2018). Neurons within the same class or subclass exhibit similar morphoelectric properties (Tremblay, Lee, and Rudy 2016; Gouwens et al. 2020), which should minimize biological variation and allow for a more appropriate technical comparison. These Cre lines are also widely used and publicly available, making them an ideal benchmark for troubleshooting for an adopting laboratory.

To evaluate success at each stage of the Patch-seq experiment, we defined a series of qualitative and quantitative parameters for each data modality (Supplementary Table 1). Although we provide our internal thresholds for each metric, they each exist along a continuum and could be applied in different ways depending on the circumstances of each individual user. Given these criteria, the protocol described here has a pass rate of 97%, 93% and 46% for electrophysiology, transcriptomics, and morphology, respectively (Figure 1), with a final rate of successful triple modality, MET, data collection of 39%. With morphological recovery having the highest rate of attrition, Patch-seq cells that fail at this point but pass electrophysiology and transcriptomics (91% in total) can still be used for spatial/anatomical, electrophysiological, and transcriptomic characterization and classification.

**Figure 1.**
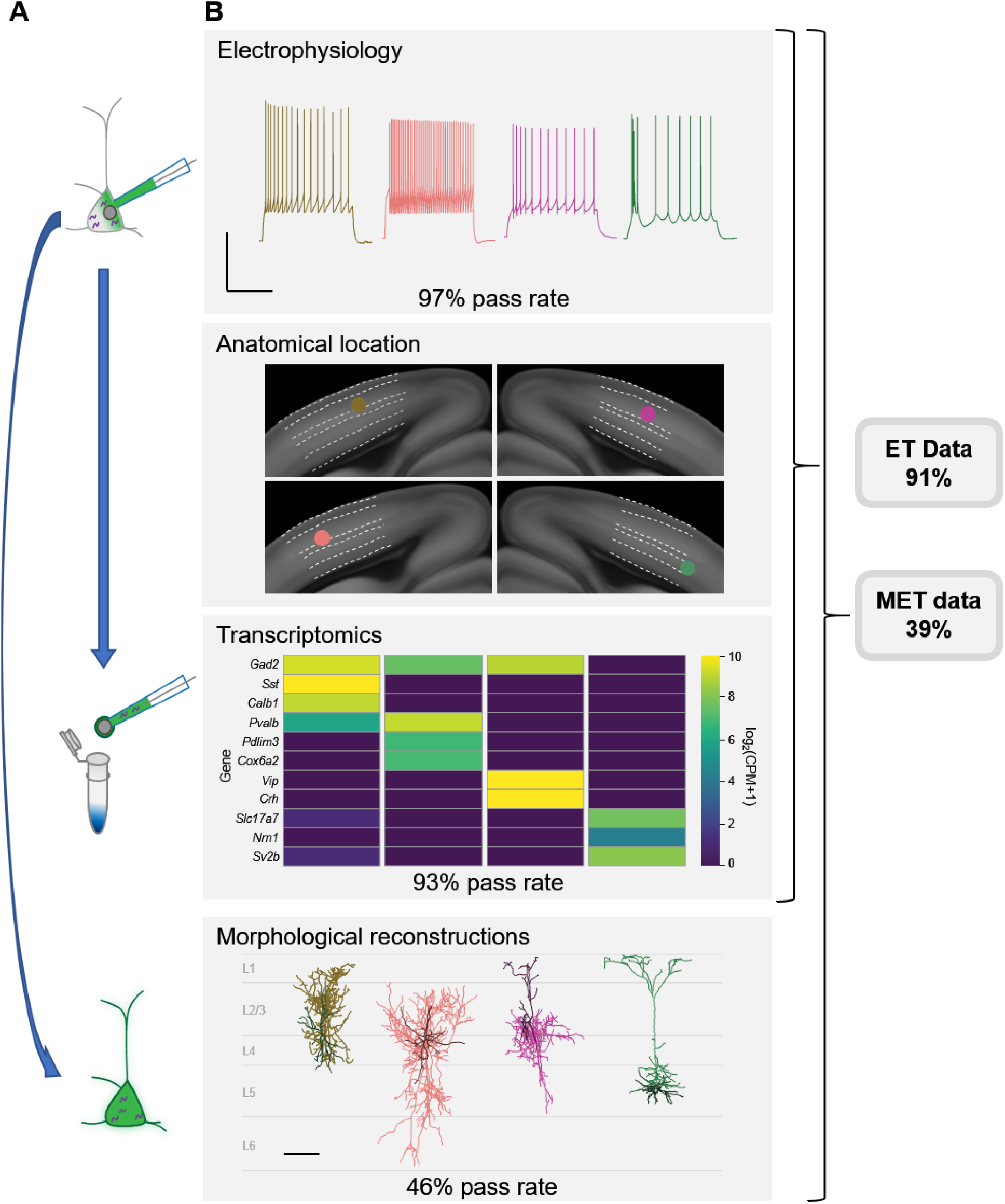
Patch-seq is a powerful technique that allows for the characterization of a cell based on electrophysiology (E), transcriptomics (T), and morphology (M). (**A**) A depiction of the sequence of events. A cell is patched to characterize the intrinsic properties of a neuron while biocytin and Alexa diffuses throughout the soma, dendrites, and axon. At the conclusion of the recording the nucleus is extracted and submitted for RNA-sequencing while the slice is fixed, stained, mounted, and assessed for morphological recovery. (**B**) Exemplar single cell examples in which we obtained all three data modalities. From left to right are inhibitory neurons from an Sst-Cre, Pvalb-Cre, Vip-Cre and an excitatory neuron from an Rbp4-Cre. Electrophysiology data shown are the voltage responses to a suprathreshold stimulus to assess a cell’s intrinsic firing properties. Scale bar is 50 mV and 500 ms. Mapping of patched cell is referenced back to a common coordinate framework to define anatomical location. Transcriptomic data are shown as a heat map of selected ‘on’ marker genes detected from the patched cell. Morphological reconstructions from the patched cells are shown, as well as their placement within the cortical layer. Data and rates represented here are from our steady-state, fully functional pipeline. Scale bar is 50 mV/500 ms for electrophysiology and 100 um for reconstructions. Cells that have a failing morphology outcome but pass for electrophysiology and transcriptomics are salvageable for ET characterization, while cells that pass all data modalities proceed for MET characterization

The patch clamp portion of the protocol consists of three phases: recording (1 - 15 mins), nucleus retrieval (< 3 mins), and nucleus extraction (~8 mins) (Figure 2). The recording phase consists of whole-cell electrophysiology to acquire intrinsic features using custom, free, publicly available software: MIES (Supplementary Figures 1, 2). The recording period is kept as short as possible to increase throughput and reduce the effect of progressive cell swelling (due to the addition of an RNAse inhibitor to the internal solution, which raises the osmolarity). Upon conclusion of the recording phase, negative pressure is applied, and the stability of the somatic membrane is monitored using visual and electrophysiological feedback.

**Figure 2.**
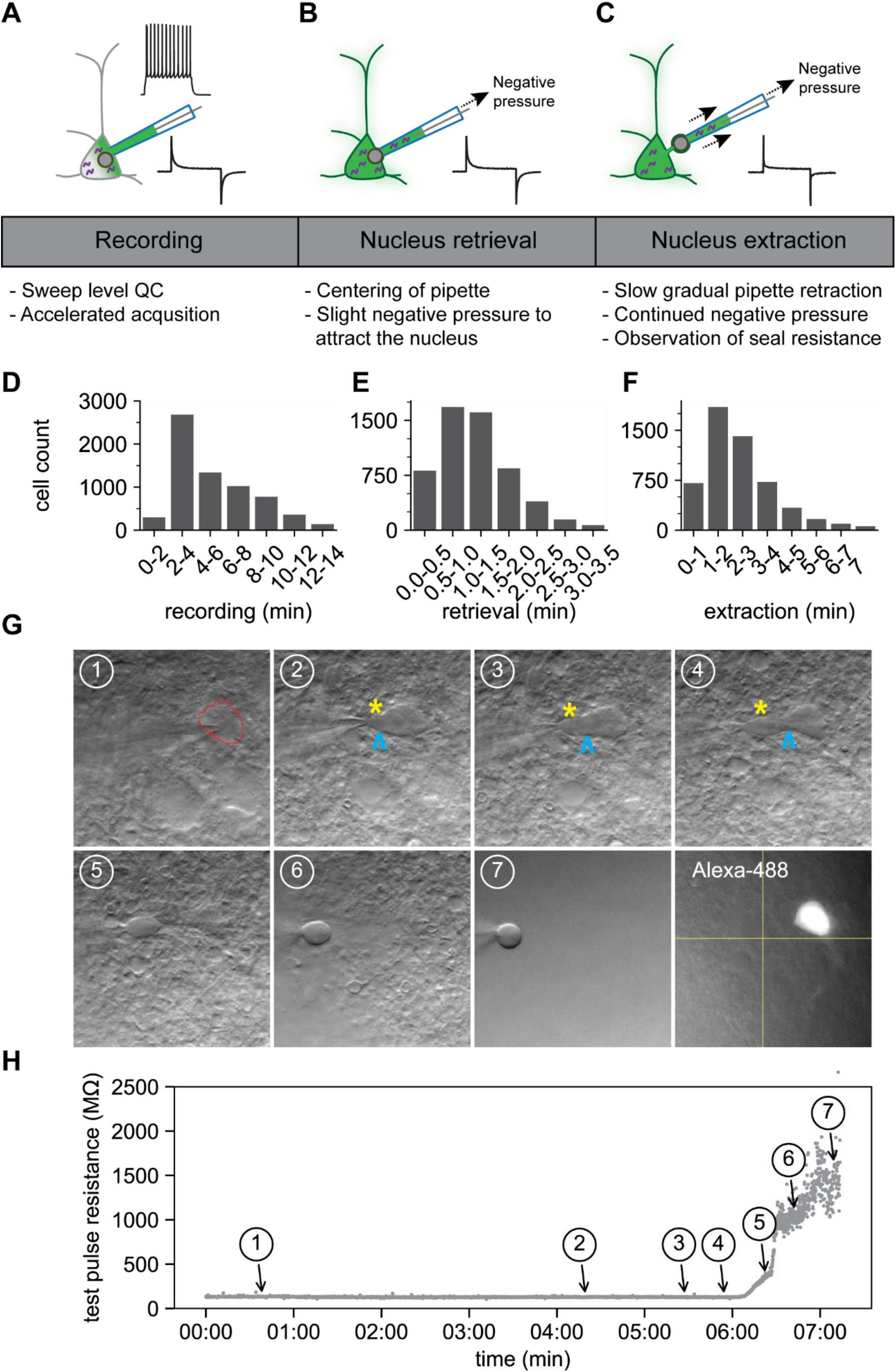
The optimized Patch-seq protocol consists of electrophysiological recording followed by nucleus retrieval and extraction. A schematic of the three major stages of the Patch-seq protocol: (**A**) recording, (**B**) nucleus retrieval, and (**C**) nucleus extraction. Histograms represent the binned time spent for (**D**) recording (N=6,720), (**E**) nucleus retrieval (N=5,692), and (**F**) nucleus extraction (N=5,442). (**G**) Time series of high-resolution images showing the gradual pipette retraction with subsequent nucleus extraction. In (**G1**), the red line denotes outline of soma and the last panel is the fill of the soma and processes visualized by Alexa-488. Yellow asterisk identifies the nucleus as it is extracted from the soma. Blue caret identifies the somatic membrane as it is stretched with the extraction of the nucleus. (**H**) is the time plot of steady state resistance, as measured from the test pulse, during the nucleus extraction phase for the cell in (**G**) with numbers corresponding to brightfield image.

The nucleus retrieval stage is focused on attracting the nucleus to the tip of the pipette by using negative pressure and moving the electrode to the location of the nucleus near center of the soma. Patience is key at this point and success relies primarily on visual feedback since the electrode resistance tends to be stable during this stage (Figure 2H, time point 1). It is important to maintain constant negative pressure during the transition from the nucleus retrieval to nucleus extraction stage.

The nucleus extraction phase requires slow retraction of the pipette along the same axis of the electrode while maintaining negative pressure. As the nucleus is pulled further from the soma, the cell membrane stretches around the nucleus and ultimately breaks, forming distinct seals around the nucleus (a ‘nucleated’ patch) and the soma (Sather et al. 1992). During this phase, constant monitoring of the membrane seal resistance is critical; the seal formation is observed electrically as a rapid rise in resistance, ideally above 1 GΩ, and referred to as end pipette resistance (endR). It is important that this stage is performed slowly and methodically; achieving the seal can take several minutes (Figures 2F, 2H). Figures 2G and 2H are time series of images and the corresponding test pulse resistance which illustrate the slow pipette retraction with an attached nucleus. The membrane finally breaks and seals between panels 5 and 7, as noted by the sharp rise in resistance shown in Figure 2H. After the nucleus has been deposited for RNA sequencing, the fluorescent Alexa dye is viewed conducted to determine if the recorded cell retained the biocytin fill (last panel in Figure 2G).

### Optimizing electrophysiology data quality and throughput

Understanding the intrinsic electrical properties of cortical neurons is a critical component of describing neuronal cell types (Gouwens et al. 2019; Gouwens et al. 2020; Scala et al. 2020; “Petilla Terminology: Nomenclature of Features of GABAergic Interneurons of the Cerebral Cortex” 2008; Markram et al. 2015; Zeng and Sanes 2017; Tremblay, Lee, and Rudy 2016). A combination of ramp and square step current injection stimulus profiles is effective at revealing discrete electrophysiological neuron types, as well as the continuum of properties between related types (Gouwens 2019). The small differences in properties between types underscores the need for 1) a large dataset to separate related groups, and 2) consistent high-quality data. We used the data from this study, now found in the Allen Cell Types Database, as a foundation to modify the custom data acquisition system to accommodate online analysis to increase the speed of data acquisition and data quality.

To facilitate flexible patch clamp data acquisition, we developed MIES, a package built on top of the Igor Pro platform. MIES has the analysis framework with automated online analysis where user code can carry out actions at predefined events. Originally designed to facilitate multichannel synaptic physiology experiments, which require easy multitasking to be performed at scale (Seeman et al. 2018). We utilized these functions to increase throughput and quality of individual Patch-seq experiments.

Patch clamp electrophysiology is traditionally based on an episodic stimulation paradigm wherein a series of ‘sweeps’ (typically < 1 s, but can be > 30 s) are acquired, in between periods of rest. Our quality control process focused on asking two fundamental questions about each sweep: 1) is the baseline prior to stimulus application stable - within and across experiments? and 2) does the cell’s membrane potential return to ’baseline’ following stimulus application? Only sweeps where both of the above conditions are true will be considered for data analysis. Although we can manually exclude data where acquisition during the stimulus indicates an issue, we find that most problems can be identified using these 2 criteria.

To avoid acquiring a sweep with an unstable baseline, a 500 ms prestimulus window to verify the resting membrane potential (RMP) was within 1 mV of the target membrane potential determined at the beginning of the experiment. To determine the stability of the RMP, we also use a threshold for the root mean square (RMS) noise of the RMP during the 500 ms baseline evaluation period. If the sweep fails either measure, it is terminated, the bias current is automatically adjusted if applicable, and the sweep is initiated again. If the number of passing sweeps required to pass the stimulus set, plus the number of failed sweeps exceeds the number of sweeps in the stimulus set, acquisition is terminated, the stimulus set fails and the user is prompted. Only once the baseline evaluation passes does the stimulus set proceed as designed (Figures 3A, 3B).

**Figure 3.**
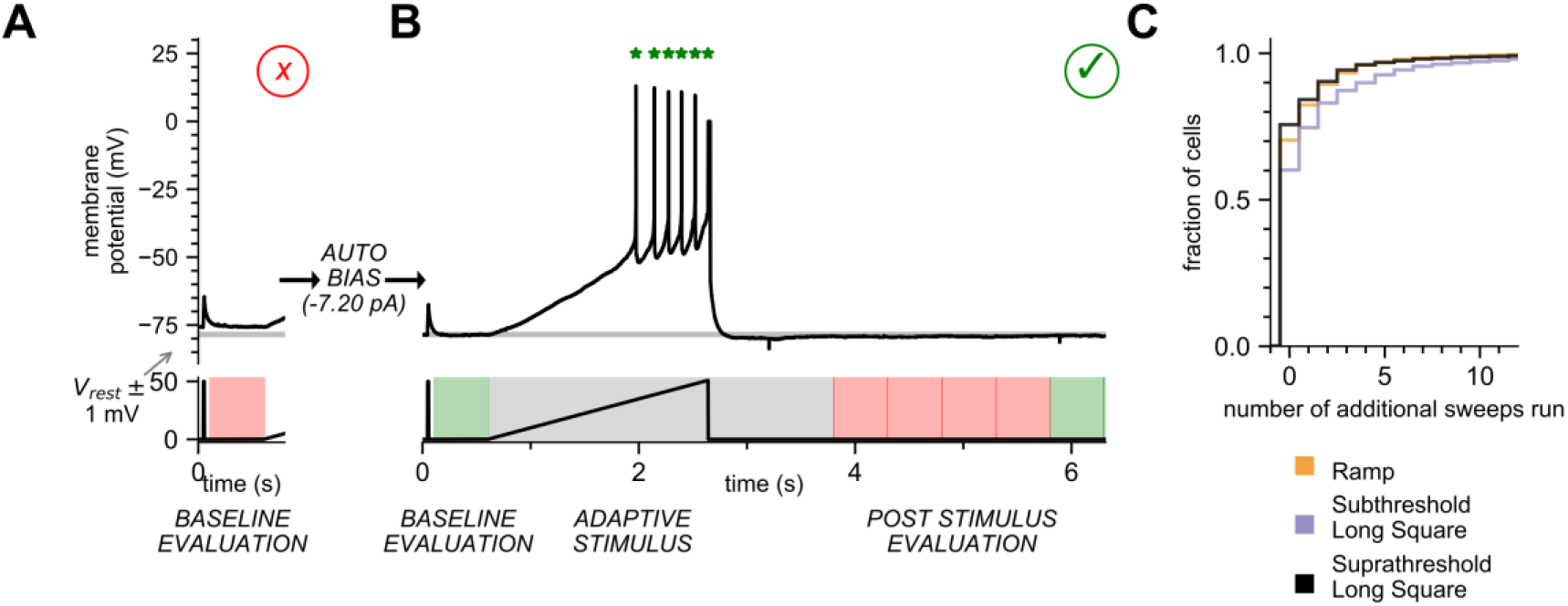
Online analysis during electrophysiology recording allows for fast, high quality data generation. (**A**) Baseline Evaluation failed. Resting Membrane Potential (RMP) was not maintained at ±1 mV (Allen institute criteria) of the established RMP (gray line at −78 mV); sweep failed and was terminated. Additional autobias current of −7.20 pA was applied during the inter-sweep interval to bring the RMP to the target value of −78 mV. (**B**) Baseline Evaluation passes and the ramp current stimulus is applied (Adaptive Stimulus). Current injection rises at 25 pA/s until 5 action potentials are detected, at which point current injection is automatically terminated. Upon conclusion of stimulus there is a 1 second minimal recovery period, followed by a assay of RMP every 500 ms (Post Stimulus Evaluation) to test for return to the initial RMP. Once the RMP recovers to within ±1 mV (Allen Institute criteria) for 2 consecutive checks acquisition is terminated and sweep is considered as ‘passed.’ (**C**) Cumulative probability plot showing the number of additional sweeps needed to qualify as passed for each of the three stimulus sets.

For each sweep, we also ensured that the post-stimulus RMP recovered to baseline (within 1 mV of target voltage) level. The time period for this ‘recovery’ can vary between cell types and can depend on the intensity of the stimulus so a ‘one size fits all’ approach, designed to accommodate the longest possible recovery time, is inefficient. To compensate, we designed a recovery period analysis function that would extend to 10 s but shortened once the target RMP was reached. Immediately after the stimulus ends, there is an absolute 500 ms minimal recovery period, followed by a continuous RMP assay in 500 ms evaluation periods. Once 2 sequential evaluation periods detect a recovered baseline, the sweep is completed and considered ‘Passed,’ ultimately completing this sweep in the minimal amount of time (Figure 3B). If two sequential periods do not pass before the end of the sweep, the sweep fails quality control (QC) and is repeated, or the user is prompted. In most instances, a stimulus set runs without triggering the repeat of a sweep (Figure 3C). However, a substantial fraction of sweeps in a stimulus set do fail, typically less than three. If these sweeps were identified as failing after the conclusion of the experiment, key sweeps may be lost. By repeating those sweeps during acquisition, the online analysis functions in MIES maximizes the chance of a successful and complete electrophysiology experiment.

One important analysis function is the automated detection of action potentials during a ramp stimulus (Figure 3B). The analysis function truncates the ramp stimulus after a predefined number (5) of action potentials have been measured, thus increasing efficiency by allowing for more standardized neuronal response and efficient data collection. Without the analysis function, the ramp stimulus would continue to depolarize the neuron regardless of when it reached the spiking threshold. Unabated depolarization may change the state, potentially fatal, of the neuron thereby altering subsequent measurements. By limiting the number of action potentials induced by the ramp, we ensure robust and consistent measurement of neuronal intrinsic features.

By default, MIES is packaged with the analysis functions used here, and elsewhere, to map the electrophysiological profile of human and mouse cortical neurons (Berg 2020; Gouwens et al. 2019; Seeman et al. 2018; Gouwens et al. 2020). These stimuli are linked such that a single click generates the entire dataset in < 3 minutes. The data can be saved directly from MIES as an NWB:N 2.0 (Ruebel et al. 2019) file, an emerging, accessible data standard for neurophysiology.

### Extracting the nucleus is key to optimal transcriptomic data quality

The mRNAs from Patch-seq extractions are often of variable quality, with some samples showing little to no content detected, whereas other samples can match or even exceed the amount of mRNA detected compared to cellular-dissociation based scRNA-seq (Tripathy et al. 2018). This variability necessitates the use of a robust, quantitative, and automated measure of mRNA quality that is independent of cell type and that can be used as an indicator for lower quality cells. To address this issue, we utilized a published methodology for assessing transcriptome quality in Patch-seq data sets (Tripathy et al. 2018). The premise of this method is that gene expression patterns of cells from matched fluorescence-activated cell sorting (FACS)-based data sets can serve as ground truth profiles for comparison to Patch-seq cells, where Patch-seq cells with technical issues are more likely to diverge from these patterns.

Three metrics are presented for assessing quality, which all rely on defining marker genes for broad cell classes of interest (‘on’ markers; e.g., Paravalbumin+ interneurons), as well as for cell types that may indicate mRNA contamination from adjacent cell bodies (‘off’ markers; e.g., astrocytes). First, the normalized marker sum (or NMS) score is a ratio of the average expression of ‘on’ marker genes for a Patch-seq cell relative to the same median expression of these genes in the matched FACS data set for the cell class with highest marker expression. This metric measures the extent to which expected genes of at least one class are expressed. Second, the contamination score assesses off-target contamination by taking the summed NMS score of all broad cell types (except the assigned class). Higher values of this metric indicate a higher likelihood that mRNA measured from a single cell also includes RNA from adjacent cells. Finally, the “quality score” is a metric aimed at capturing both types of technical issues by measuring the correlation of ‘on’ and ‘off’ marker genes in Patch-seq cells with the average expression profile of dissociated cells of the same type.

Here we expand on the work of Tripathy et al. by defining marker genes for class using a more recent study of single cells collected from mouse primary visual cortex (VISp) and anterolateral motor cortex (ALM) (Tasic et al. 2018). We chose 50 marker genes for each class, defining ‘on’ markers by subclass, and ‘off’ markers by subclass for non-neuronal cells and by class for neuronal cells. Figure 4A displays gene expression data (counts per million and log2-transformed) and a corresponding high NMS score, from cells targeted by specific Cre lines and their representative ‘on’ marker genes. We find that the NMS distribution is roughly bimodal (Figure 4A) and we have chosen 0.4 as a rough cut-off for high- and low-quality data (Gouwens et al. 2020). Additionally, a low NMS score is more reflective of a lack of detectable genes and not necessarily an increase in ‘off’ marker expression (genes highlighted in gray in Figure 4A). We sought to investigate the relationship of the ‘on’ marker genes of an assigned subclass and how they relate to cells that were patched from a matching Cre line. Cells with a higher NMS score from a Cre line are generally assigned to the appropriate subclass, whereas those with a low NMS score have more promiscuous assignments (Figure 4B).

**Figure 4.**
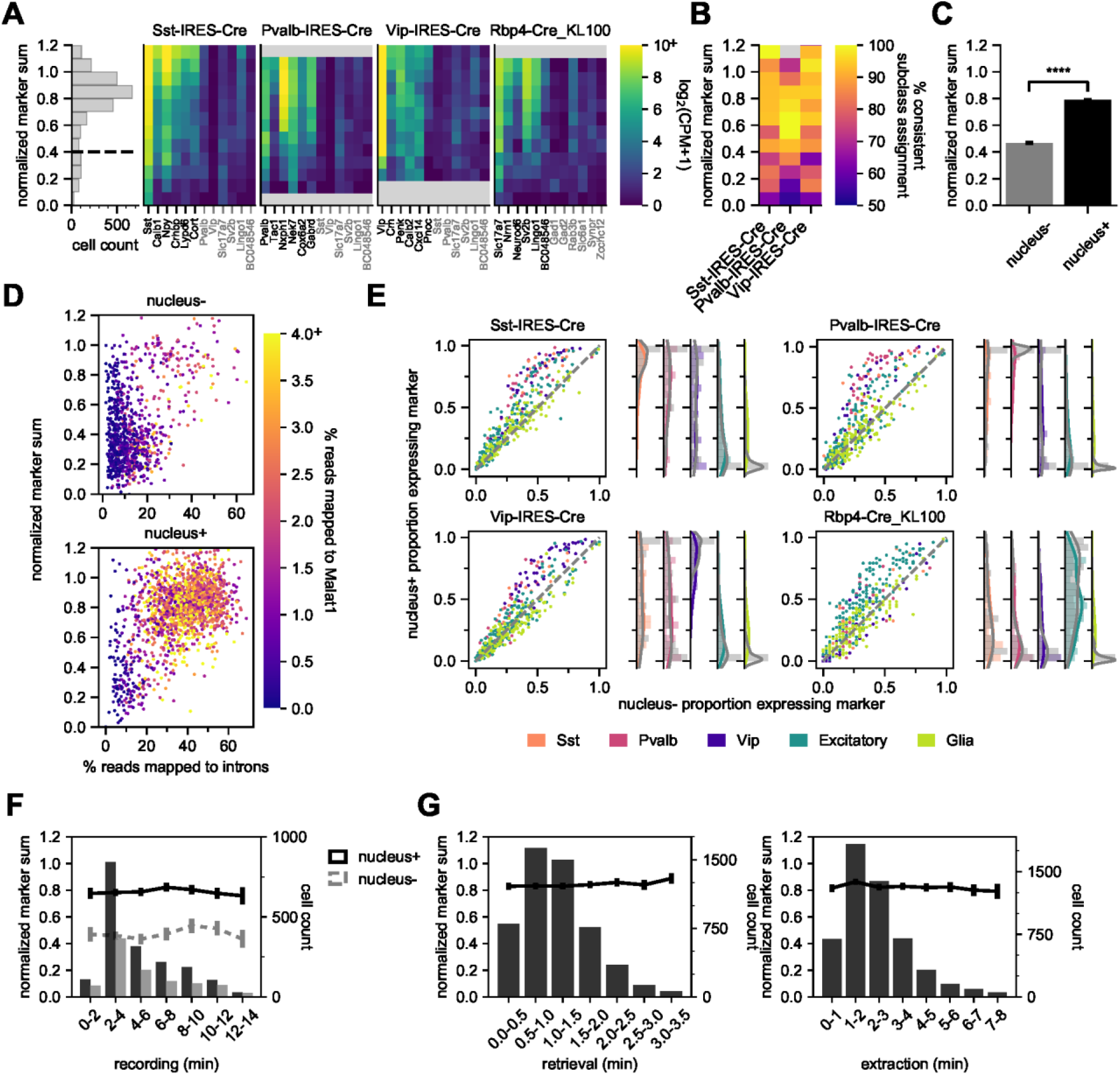
Nucleus extraction is key to transcriptomic success. To examine the quality of Patch-seq samples we used the NMS score. The first panel (**A**) is a histogram that displays the range of scores from all cells from three of the major inhibitory Cre lines (Sst-Cre, Pvalb-Cre, and Vip-Cre) and one excitatory Cre line (Rbp4-Cre) with the dotted line, at 0.4, indicating the separation between pass and fail (N=2,717). The heat maps display the average gene expression of ’on-marker’ (in black) or ‘off-marker’ (in gray) genes for these cells. Gene expression values are log2(CPM+1). (**B**) A heatmap showing how the NMS score relates to the % of genes assigned to a specific subclass and the corresponding Cre line. **(C)(D)(F)** examine an early dataset of Patch-seq samples with both nucleus+ and nucleus-outcomes (N=1,804 nucleus+ samples; N=885 nucleus-samples). The bar plots in (**C**) represent the difference in NMS scores. (**D**) A scatter plot viewing the distribution of % reads mapped to introns and how they related to the NMS score for nucleus+ (top) and nucleus- (bottom). Color ramp indicates the % reads mapped to Malat1. (**E**) Scatter plots examining the relationship nucleus+ or nucleus- and the presence of subclass-specific marker gene expressions for 4 different Cre lines. The colors of dots represent the subclass to which the gene is specific (N=50 Sst, N=50 Pvalb, N=50 Vip, N=193 Excitatory, N=146 Glia genes). The histograms are the quantification of subclass-specific genes for each of the respective Cre lines for nucleus+ samples only. The gray bars are FACS data. In (**F**), the histograms represent the binned time spent and the cell count for recording duration, the darker bars plot successful extraction of the nucleus (nucleus+) whereas the lighter bars plot failed extraction of nucleus (nucleus-). Solid line represents the NMS score for nucleus+ samples as a function of time, whereas gray dotted line represents nucleus-samples. (**G**) represents the binned time spent and cell count for nucleus retrieval and nucleus extraction (N=5,230). The solid line represents the NMS score as a function of time.

Using the marker gene list and calculated NMS score to evaluate transcriptomic quality, we have determined that Patch-seq experiments where we collect the cytosol and nucleus (nucleus+) have significantly higher NMS scores than cytosol-only (nucleus-) samples, *t*(2,687)=32.3, *p*<0.0001 (Figure 4C). We have designed two sets of metrics to evaluate and confirm the presence of the nucleus in Patch-seq samples: nuclear and subclass-specific gene expression. Gene expression data is a combined measurement of intronic reads (which are localized to the nucleus) and exonic reads (which are found throughout the soma); therefore, nucleus+ samples will have a higher percentage of intronic reads (Gaidatzis et al. 2015). Additionally, we examined the presence of Metastasis Associated Lung Adenocarcinoma Transcript 1 (*Malat1*), which has been found as a mammalian-specific nucleus-related gene (Hutchinson et al. 2007). Both the intronic reads and *Malat1* gene expression are correlated with the fraction of RNA collected from the nucleus relative to the cytoplasm (Bakken et al. 2018). Indeed, with our nucleus+ Patch-seq samples we observed a higher correlation with *Malat1* expression and the percentage of reads mapped to introns compared to nucleus-samples (Figure 4D). These measures demonstrate that nucleus+ samples have higher transcriptomic quality but can also guide future experiments as confirmatory evidence for nuclei collection.

In addition to intronic reads and nucleus-specific genes, nucleus+ samples have a higher detection of overall genes relative to nucleus-samples (Supplementary Figure 3A). More specifically the nucleus+ samples have a higher fraction of ‘on’ marker gene expression specific to the appropriate Cre lines. Whereas ‘off’ marker genes, genes not specific to the Cre line, or glia-related genes, are less differentiated between nucleus+ and nucleus- samples (Figure 4E). The histograms to the right of each scatter plot quantify the average subclass-specific gene expression for nucleus+ samples and how they relate to their FACS (gray) counterparts. One important item to note is the higher detection of glia-related genes in Patch-seq samples compared to FACS. This is expected due to the inherent nature of the Patch-seq process as the patch pipette navigates through the tissue. We can conclude from these plots that the detection of subclass-specific gene expression in nucleus+ samples is similar to that of FACS-isolated samples.

Due to the sensitivity and unknown stability of the neuron in the dialyzed whole-cell configuration, we sought to determine if there was a relationship between the experiment duration and the quality of extracted mRNA and subsequent cDNA (Sucher et al. 2000; Veys et al. 2012). To do this we tracked the time spent at each one of the Patch-seq stages (Figures 4F, 4G). Using the NMS score as a measure of quality, surprisingly, we found no effect of time on the quality of the transcriptomic content. As shown in Figures 4C and 4F, nucleus+ conditions have a significantly higher NMS score with no effect of patch duration. Additionally, the time of nucleus retrieval and extraction phases did not affect the NMS score. This corroborates other findings (Cadwell et al. 2017) that the integrity of the RNA is preserved during the Patch-seq recordings. It is noteworthy and informative that the range of extraction times can take less than 1 minute or up to 8 minutes to successfully extract the nucleus. Despite the lack of an effect, it may still be advisable to keep the duration shorter for other reasons such as to allow for higher throughput or for optimization of morphological recovery.

Additionally, the cDNA can be evaluated to inform the quality of the Patch-seq samples. Electrophoretograms are obtained from a fragment or bioanalyzer and can provide unique metrics about the amplified cDNA. Quantitatively, two parameters can be obtained: 1) the amount of amplifiable content, and 2) the quality of the cDNA, measured as the ratio of high base pair material. Qualitative analysis can provide confirmation about the quantitative data obtained, such as proper distributions and shape of the electrophoretogram, positive controls, and artifacts. We found the NMS score is highly correlated with each metric and a clear separation between nucleus+ and nucleus- samples. The difference between the two outcomes (nucleus+ vs. nucleus-) was found to be highly significant for each metric: cDNA quantity, t=14.91, p<0.001 and cDNA quality, t=22.77, p<0.001 (Supplementary Figures 3B, 3C). Receiver operating characteristic (ROC) curve analyses were performed to evaluate the sensitivity and specificity of the cDNA for the presence or absence of the nucleus in Patch-seq samples. As shown in Supplementary Figure 3B and 3C, both the quantity and quality of the cDNA achieved an area under the curve (AUC) was >0.80, demonstrating the presence of the nucleus is strongly correlated with high quality data.

### Optimizing morphology success with Patch-seq recordings

The shape and morphological features of a neuron are important for its functional output and can be used to classify and define types (Zeng and Sanes 2017; Gouwens et al. 2019; Harris and Shepherd 2015; Markram et al. 2015; Lodato and Arlotta 2015). Obtaining morphologies from Patch-seq samples has been problematic and strategies for improving recovery have not been thoroughly examined. Early Patch-seq studies were unable to recover the morphology of the patched neuron and had to rely on electrophysiological classifiers to infer the morphological properties (Fuzik et al. 2015; Cadwell et al. 2015). More recently, (Cadwell et al. 2017) has described success in recovering morphologies from Patch-seq samples using longer recording times but RNA quality in these cells was generally lower than reported for FACS-isolated cells. We have shown that extracting the nucleus (nucleated patches) are key in transcriptomic success; historically, this paradigm has been used for studies of membrane biophysics. In previous studies using nucleated patches for this purpose (Eyal et al. 2016; Bekkers 2000; Gurkiewicz and Korngreen 2006), morphological recovery of the neuron has not been well studied. Here we standardized metrics measuring the neuron morphology outcomes, then used those evaluations to adjust the cell recording protocol to optimize morphological recovery while retaining high-quality RNA extraction.

Figures 5A-5E displays representative examples of morphological outcomes and evaluations with the Patch-seq technique. ‘High quality’ fills have a visible soma and processes and are suitable for 3D digital reconstructions as shown in Figures 5A and 5B for an excitatory and inhibitory neuron, respectively. ‘Insufficient axon’ fills represent cells with filled dendrites, but weakly filled axons or axons that are orthogonal and exit the slice (Figure 5C). ‘Medium quality’ fills have a visible soma and some processes but are not suitable for 3D digital reconstruction (Figure 5D). ‘Failed fills’ have no visible somas and likely result from the collapse of the soma during nucleus extraction and subsequent leakage of the biocytin (Figure 5E). A single coronal slice with multiple Patch-seq recordings can lead to varying outcomes for morphological quality (Supplementary Figure 4), suggesting that factors prior to the slice processing phase are critical determinants of morphological recovery. Here we focused on the impact of recording variables on morphological recovery outcomes.

**Figure 5.**
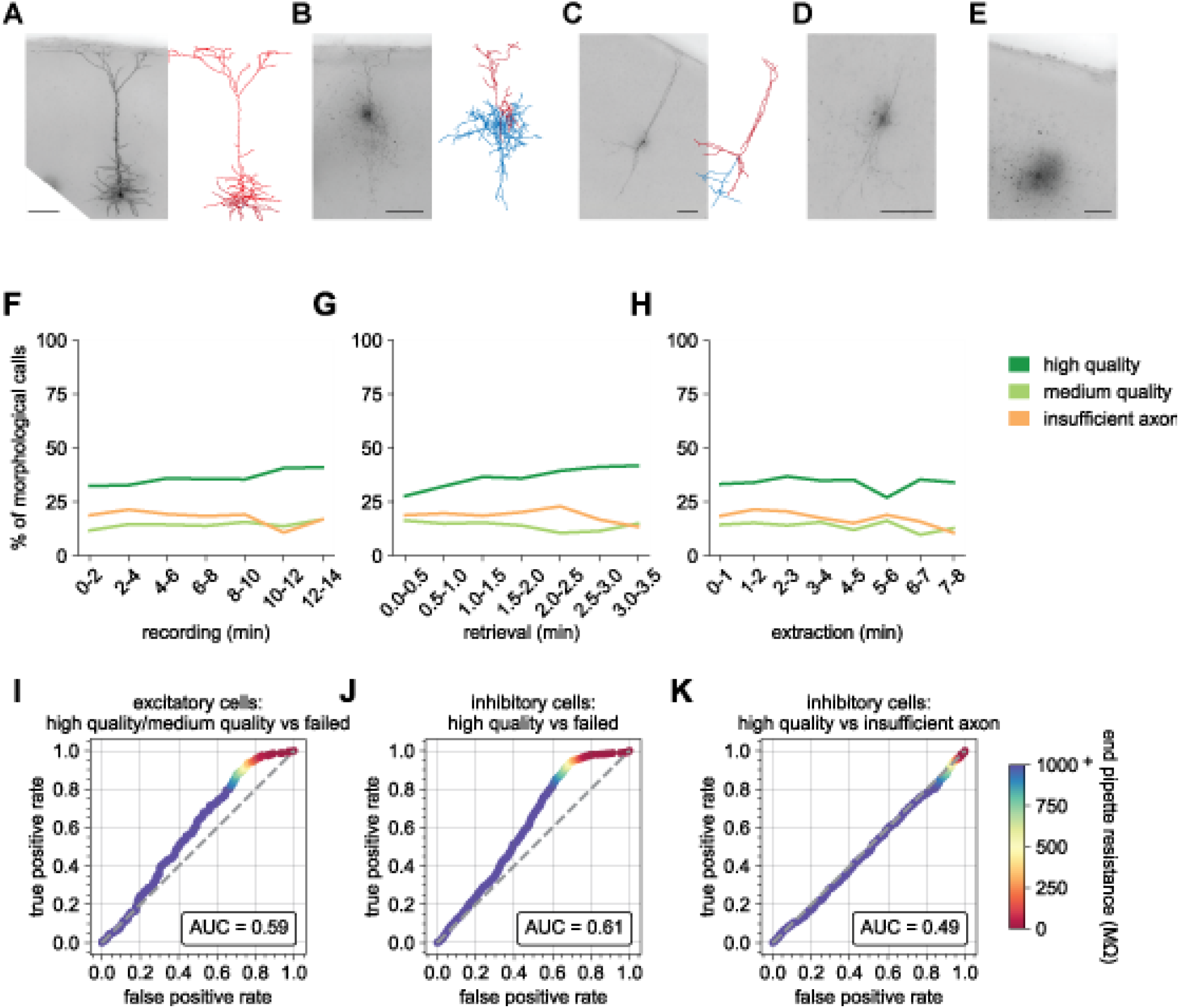
End pipette resistance, but not recording duration, is a significant predictor of morphology success. Example biocytin recovery and subsequent morphological reconstructions of a high quality filled Rbp4-Cre+ excitatory (**A**) and a Vip-Cre+ inhibitory (**B**) neuron. (**C**) A Vip-Cre+ inhibitory neuron that failed due to an insufficiently filled axon. (**D**) An interneuron that was weakly filled and was classified as medium quality. (**E**) A failed fill where no processes are visible and biocytin leakage is apparent. (A-E) are high-resolution 63x stack minimum image projection (MIP) images and (**A-C**) have the corresponding morphological reconstruction with dendrites in red and axon in blue. Line plots demonstrate the time spent for each of the Patch-seq phases: (**F**) recording (N=5,554), (**G**) nucleus retrieval (N=5,563), and (**H**) nucleus extraction (N=5,342), and how they relate to the morphological recovery. ROC analyses comparing high/medium quality versus failed morphology outcomes for patched (**I**) excitatory (N=656 high/medium quality; N=373 failed) and (**J**) inhibitory neurons (N=1,054 high quality; N=1,215 failed). (**K**) ROC analysis comparing N=1,054 high quality versus N=878 insufficient axon morphology outcomes for patched inhibitory neurons. Heat map of the ROC curve is the end pipette resistance (MΩ) measured at the conclusion of nucleus extraction.

Many studies have shown that patch duration must range from 15 mins up to 60 mins to obtain optimal filling of neuronal processes with biocytin (Gouwens et al. 2019; Marx et al. 2012; Cadwell et al. 2017). To maximize throughput, we targeted a recording duration < 10 mins. Surprisingly, we found no effect of the recording duration (range: 3 to 12 minutes) on the fraction of cells that were deemed optimal for morphology reconstruction (~45%, Figure 5F). The duration dependence of morphology output was mostly flat for other phases of the recording process (Figures 5G, 5H), with perhaps a slight upward trend in outcome for longer retrieval times (Figure 5G). To investigate the effect that recording multiple neurons per slice has on morphology outcomes, we compared the morphologies of neurons from brain slices that had 1, 2, 3, or 4 recordings before slice fixation. We found that multiple Patch-seq recordings could be obtained in a single slice with no deleterious effects as the prior patched cell(s) remained quiescent during subsequent recordings. We did observe a trend in which the last cell recorded in the slice had the poorest outcome for ‘high quality’ with an increase in ‘insufficient axon’ or ‘failed’ outcomes (Supplementary Figure 5A), indicating the possibility of insufficient time for complete diffusion of biocytin throughout the neuron prior to fixation. This is consistent with a previous report (Cadwell et al. 2017) showing that when the slice is fixed immediately after a single neuron is recorded, longer recording durations are required for sufficient fill.

We next asked how the resistance of the nucleated patch membrane (end pipette resistance, endR, Figure 2H) predicts the ultimate cell morphology, as it likely reflects how well the membrane reseals around both the extracted nucleus and the neuron remaining in the slice. We found the endR to be highly predictive of the final morphology outcome for both excitatory and inhibitory neurons. When comparing the outcome of high and medium quality fills (combined) versus failed fills using a receiver operating characteristic (ROC) curve, we find an area under the curve (AUC) of 0.57 and 0.62 for excitatory and inhibitory cells, respectively. Most interestingly, there is a prominent shoulder in the ROC curve at an endR of 100 MΩ. Using this value as a cutoff can reduce the fraction of morphology failures by about 30% at a cost of < 5% of high and medium morphologies for both excitatory and inhibitory cells (Figures 5I, 5J). EndR did not impact the insufficient filling of the cell, which is likely influenced by the recording duration or positioning of the cell within the slice (Figure 5K). This demonstrates that recordings that end with a pipette with a high endR are more successful at retaining the morphological fill.

Unsurprisingly, cell health is also a significant factor in the ability to retain the morphology of the recorded neuron. We performed a qualitative rating on a scale of 1 (worst) to 5 (best) of cell health. This rating included a visual assessment, using metrics such as soma shape and sharpness of the plasma membrane, and a recording quality assessment, using metrics such as baseline stability and number of failed sweeps. We found that cells ranked 1 or 2, had a lower chance of a ‘high quality’ score than cells with rank of 3, 4 or 5. Interestingly, there was little to no improvement of ‘high quality’ fills between rankings of 3 and 5 (Supplementary Figure 5B). Supplementary Figure 6 shows representative 63x resolution minimum image projections (MIPs) of biocytin fills and their resultant morphological calls compared to the cell health assessment score.

### Application to other species and cell types

A powerful aspect of the FACS-based transcriptomic studies is the ability to use similar methods to profile and compare neurons from different brain regions or species (Bakken et al. 2018; Hodge et al. 2019; Kalmbach et al. 2020). With that in mind, we co-developed this protocol in both mouse tissue and tissue from human surgical resections. We sampled 1,402 human neocortical neurons across the four lobes (Supplementary Figure 7A) and observed a pass rate of 76%, 77% and 47% for electrophysiology, transcriptomics and morphology, respectively. Ultimately, we have final rate of 69% for ET and 28% for MET characterization (Supplementary Figure 7B) – all of which are comparable to mouse rates of success. Importantly, the key predictors of success – collecting the nucleus for transcriptomics and end electrode resistance for morphology, were consistent between mouse and human experiments (Supplementary Figure 7C, 7D). In one example, we recorded from 10 cells in the same human cortical slice (Figure 6) – for each neuron, the nucleus was extracted and the end electrode resistance was > 1 GΩ. The fact that a single human slice yields 100% success despite lower population averages highlights that higher success rates are feasible, especially in human tissue, which has a number of experimental variables. Gene expression was consistent with the mapped transcriptomic types (Figure 6C) and morphoelectric properties of each cell are consistent with the cell types to which they are mapped. Glutamatergic neurons are pyramidal and have regular, adapting spiking patterns, while the GABAergic neurons show a greater diversity in their morphologies and firing patterns. To further demonstrate the generalizability of the approach, we targeted pyramidal neurons in Macaque *ex vivo* acute brain slices from the temporal cortex region. We collected high-quality, morphoelectric and transcriptomic data from 36 neurons that show that quality metrics were similarly correlated with nucleus extraction (Supplementary Figure 8) as shown with mouse and human Patch-seq experiments.

**Figure 6.**
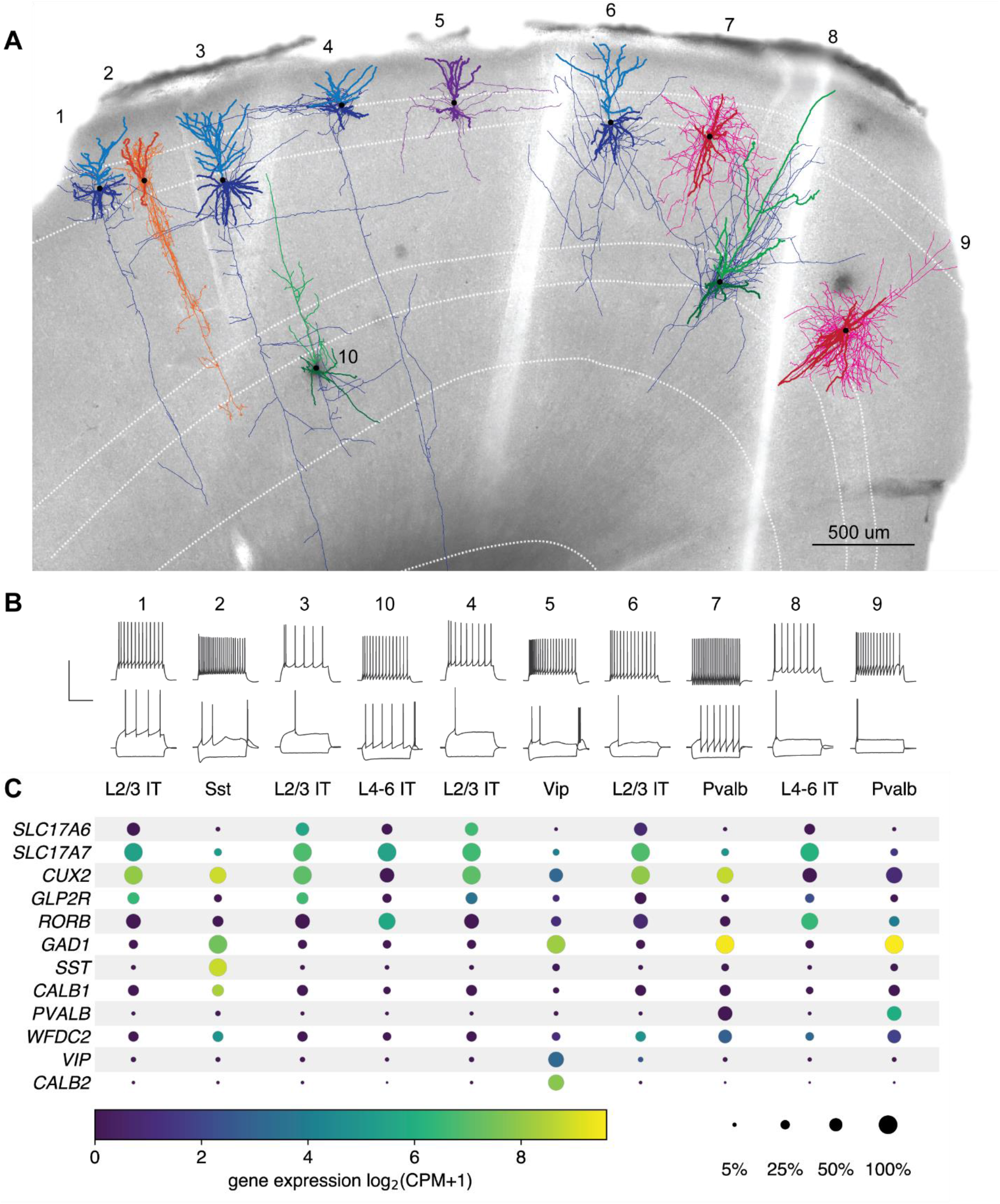
Exemplar human neocortical slice with 10 successful Patch-seq recordings. (**A**) is a 20x bright field image of a human acute neocortical slice containing biocytin reacted fills with overlaid digital reconstructions from ten Patch-seq recordings. Reconstructions are colored by their corresponding mapped subclass: L2/3 IT - blue, L4-6 IT - green, Sst - orange, Vp - purple and Pvalb-pink. Axons are light-weighted lines and the basal/apical dendrites for excitatory neurons are distinguishable by darker and lighter colors, respectively. Cells are numbered in the order in which they were patched. Scale bar is 500 um. (**B**) Voltage responses of corresponding cell to 1 s-long current steps equal to −70 pA and rheobase (bottom traces) and rheobase + 80 pA (top trace). Scale bar, vertical 80 mV, horizontal 500 ms. (**C**) Dot plot of marker genes that are differentially expressed across human cortical neurons. Expression values (color) for each cell in slice pictured in (**A**) are log-transformed; size of dot indicates percentage of cells with non-zero expression of that gene within the corresponding subclasses in dissociated cells from FACS. Percentages calculated from N=3,706 L2/3 IT; N=5,636 L4-6 IT; N=809 Sst; N=1,298 Vip; N=809 Pvalb dissociated cells.

## Discussion

We have optimized the Patch-seq technique using a standardized approach to provide a comprehensive protocol and guidance for others in the community. Our findings build upon key components of previous detailed Patch-seq protocols (Cadwell et al. 2017; Fuzik et al. 2015) to increase throughput using custom software and identify key predictors of data quality, most critically extracting the nucleus and maintaining membrane integrity as the electrode is retracted. The large size of our dataset (combined mouse and human N=8,122) has allowed us the opportunity to examine the relationship between specific QC metrics and the resultant experimental outcomes. We have provided a guide for each metric including the range and what type of data to expect. For example, due to this high cost of sequencing, it might be in the financial interest to evaluate the cDNA metrics and be judicious about which samples to send for sequencing. More importantly, these serve as indicators to guide experiments and allow the user to make predictions and to optimize the approach for their specific application.

This protocol has resulted in the successful collection of large-scale Patch-seq data from mouse and human brain cell types, and this data are publicly accessible as part of the Allen Cell Types Database (https://portal.brain-map.org) as well as the NIH’s Brain Research through Advancing Innovative Neurotechnologies (BRAIN) Initiative - Cell Census Network (BICCN, https://biccn.org). To accelerate data collection in the scientific community, we identified 2 use cases for an independent researcher may want to generate comparable data:

### 1) Determining the morphoelectric phenotype of cells corresponding to a large (FACS-based) transcriptomics dataset

The Patch-seq platform makes morphoelectric data accessible to a transcriptomics community that is accustomed to using large datasets to probe biological variance within a population. However, previous descriptions of the Patch-seq method have detailed a low-throughput technique, suitable for targeted small studies (Cadwell et al. 2017; Lipovsek, Browne, and Grubb 2020). Here we describe an approach that can be used to generate combined morphoelectric and transcriptomic data at a scale that approaches recent transcriptomic studies. Indeed, the number of Patch-seq cells in a recent manuscript using this protocol (4,270 cells, (Gouwens et al. 2019)) was more than 2.5 times the number of cells in a FACS-based single cell transcriptomics study just a few years ago (1,679 cells, (Tasic et al. 2016)).

### 2) A smaller study that ‘extends’ the Allen Cell Types database to answer a specific research question

Building upon large standardized datasets has been a successful way to address challenging questions (Campbell et al. 2017; Steinmetz et al. 2019). However, in the case of Patch-seq, combined analysis of electrophysiological data generated in different labs has proved challenging to integrate (Tripathy et al. 2018). To combat this, here we detail both our approach and any quality metrics we use to exclude data. Another advantage using the tools described here is that MIES saves data in NWB:N 2.0, the open data format adopted by a number of labs, including all electrophysiology data deposited in the Allen Cell Types Database and BICCN. Pre-saved stimulus sets and the ability to save data in a standardized format will facilitate combined analysis of new and archived data.

Despite the improvements described here, there is room for additional optimization of the Patch-seq technique, primarily in the area of morphological recovery, where ~46% of experiments result in cells fit for digital reconstruction. Since morphological recovery is related to the qualitative ranking of cell health, additional optimization of the tissue slicing protocol tailored to specific mouse age, region, or cell types (Ting et al. 2018) could lead to improved recovery. An additional area for further investigation is the effect of the size of the electrode on electrophysiological, transcriptomic, and morphological data quality. Since the seal resistance following retraction is a strong predictor of morphology outcome, it may be that smaller electrodes, which would disturb less of the total cell membrane, would be predicted to improve morphological recovery success. However, we initially found that smaller electrodes made it difficult to collect the nucleus, so we did not pursue this strategy further. Depending on the scientific question being asked, one can make different trade-offs between potentially conflicting data modalities.

To improve throughput, the patch-clamp process could be further automated (Kodandaramaiah et al. 2016). Indeed, here we have shown that automated analysis of electrophysiological features improves both the speed of acquisition and the quality of the ultimate data product. Automation could clearly improve throughput, allowing the user to focus on getting the next cell while the automated patch-clamp device initiates a recording. Implementing automation could also improve the data quality in other modalities in Patch-seq. For example, automated retraction of the pipette during nucleus extraction, with endR feedback, could result in a slower, more standardized movement with the potential to improve the rate of morphological recovery.

The protocol described here is focused on the interrogation of each cell’s intrinsic electrical properties, but an obvious extension would be to incorporate Patch-seq with measurement of syaptic properties. Characterizing the connectivity and synaptic dynamics has been shown to differentiate classes of neurons (Jiang et al. 2015; Seeman et al. 2018; Földy et al. 2016) and linking the rates or strength of synaptic connections with genes of interest and subsequent transcriptomic types would further our understanding of the functional role of cell types.

Large-scale transcriptomic studies have led to an incredible amount of progress in understanding the degrees of cell type differences between brain regions (Yao et al. 2020; Tasic et al. 2018) and species (Bakken et al. 2018; Hodge et al. 2019; Bakken et al. 2016). To probe the phenotypic consequences of this differentiation, a platform is required that is robust to these variables. We have successfully used the Patch-seq protocol described here to study non-human primate and human excitatory neurons (Kalmbach et al. 2020; Berg 2020; Bakken 2020), mouse cortical interneurons (Gouwens et al. 2020; Keaveney et al. 2020). Here we demonstrate that the key to successfully adapting the protocol to a new system, like the human or non-human primate, is to focus on consistency in a few techniques, including nucleus extraction and maintaining membrane integrity during retraction. By dividing the protocol up into clear metrics and benchmarks, implementing labs can adapt critical parts of the protocol while monitoring trade-offs in other modalities. Intriguingly, in some human cases, we are able to achieve high quality triple modality data from many cells in a single slice. By continuing to monitor experiment variables, in this case such as patient age, brain region, or underlying pathology.

Here we demonstrate that the key to successfully adapting the protocol to a new system, like the human or non-human primate, is to focus on consistency in a few techniques, including nucleus extraction and maintaining membrane integrity during retraction. Despite its success, there are likely cell types for which this protocol will not be feasible as-is, due to factors like cell size or cell excitability. By dividing the protocol up into clear metrics and benchmarks, implementing labs can adapt critical parts of the protocol while monitoring trade-offs in other modalities.

As FACS-based single cell transcriptomics establishes a foundation for studying neuronal cell types using large, standardized datasets with common tools, complementary techniques must adapt to facilitate faster means of data acquisition that can be applied to diverse tissues, and with transparent quality metrics. Patch-seq, although a relatively young technique, has shown a scalability that holds promise for accelerating progress toward understanding of the phenotypic consequence of transcriptomic variability. The protocol described here details the keys to that scalability and provides the tools to allow others to adapt the approach for diverse applications in their own research programs.

## Methods

### Mouse breeding and husbandry

All procedures were carried out in accordance with the Institutional Animal Care and Use Committee at the Allen Institute for Brain Science. Animals (<5 mice per cage) were provided food and water ad libitum and were maintained on a regular 12 hour light–dark cycle. Animals were maintained on the C57BL/6J background, and newly received or generated transgenic lines were backcrossed to C57BL/6J. Experimental animals were heterozygous for the recombinase transgenes and the reporter transgenes.

### Human tissue acquisition

Human tissue acquisition details can be found here (Berg 2020). Briefly, surgical specimens were obtained from local hospitals (Harborview Medical Center, Swedish Medical Center and University of Washington Medical Center) in collaboration with local neurosurgeons. All patients provided informed consent and experimental procedures were approved by hospital institute review boards before commencing the study. Human surgical tissue specimens were immediately transported (15-35 min) from the hospital site to the laboratory for further processing.

### Tissue Processing

For mouse experiemtns male and females were used between the ages of P45 and P70 and anesthetized with 5% isoflurane and intracardially perfused with 25 mL of ice-cold slicing artificial cerebral spinal fluid (ACSF; 0.5 mM calcium chloride (dehydrate), 25 mM D-glucose, 20 mM HEPES buffer, 10 mM magnesium sulfate, 1.25 mM sodium phosphate monobasic monohydrate, 3 mM myo-inositol, 12 mM *N*-acetyl-L-cysteine, 96 mM *N*-methyl-D-glucamine chloride (NMDG-Cl), 2.5 mM potassium chloride, 25 mM sodium bicarbonate, 5 mM sodium L-ascorbate, 3 mM sodium pyruvate, 0.01 mM taurine, and 2 mM thiourea (pH 7.3), continuously bubbled with 95% O_2_/5% CO_2_). Human, mouse or macaque slices (350 μm) were generated (Compresstome VF-300 vibrating microtome, Precisionary Instruments or VT1200S Vibratome, Leica Biosystems), with a block-face image acquired (Mako G125B PoE camera with custom integrated software) before each section to aid in registration to the common mouse reference atlas. Brains were mounted for slicing either coronally or 17° off-coronal to preserve intactness of neuronal processes in primary visual cortex.

Slices were transferred to an oxygenated and warmed (34 °C) slicing ACSF for 10 min, then transferred to room temperature holding ACSF (2 mM calcium chloride (dehydrate), 25 mM D-glucose, 20 mM HEPES buffer, 2 mM magnesium sulfate, 1.25 mM sodium phosphate monobasic monohydrate, 3 mM myo-inositol, 12.3 mM *N*-acetyl-L-cysteine, 84 mM sodium chloride, 2.5 mM potassium chloride, 25 mM sodium bicarbonate, 5 mM sodium L-ascorbate, 3 mM sodium pyruvate, 0.01 mM taurine, and 2 mM thiourea (pH 7.3), continuously bubbled with 95% O_2_/5% CO_2_) for the remainder of the day until transferred for patch-clamp recordings.

### Patch-clamp recording

Slices were bathed in warm (34 °C) recording ACSF (2 mM calcium chloride (dehydrate), 12.5 mM D-glucose, 1 mM magnesium sulfate, 1.25 mM sodium phosphate monobasic monohydrate, 2.5 mM potassium chloride, 26 mM sodium bicarbonate, and 126 mM sodium chloride (pH 7.3), continuously bubbled with 95% O_2_/5% CO_2_). The bath solution contained blockers of fast glutamatergic (1 mM kynurenic acid) and GABAergic synaptic transmission (0.1 mM picrotoxin). Thick-walled borosilicate glass (Warner Instruments, G150F-3) electrodes were manufactured (Narishige PC-10) with a resistance of 4–5 MΩ. Before recording, the electrodes were filled with ~1.0-1.5 μL of internal solution with biocytin (110 mM potassium gluconate, 10.0 mM HEPES, 0.2 mM ethylene glycol-bis (2-aminoethylether)-*N*,*N*,*N*’,*N*’-tetraacetic acid, 4 mM potassium chloride, 0.3 mM guanosine 5’-triphosphate sodium salt hydrate, 10 mM phosphocreatine disodium salt hydrate, 1 mM adenosine 5’-triphosphate magnesium salt, 20 μg/mL glycogen, 0.5U/μL RNAse inhibitor (Takara, 2313A) and 0.5% biocytin (Sigma B4261), pH 7.3). The pipette was mounted on a Multiclamp 700B amplifier headstage (Molecular Devices) fixed to a micromanipulator (PatchStar, Scientifica).

The composition of bath and internal solution as well as preparation methods were made to maximize the tissue quality of slices from adult mice, to align with solution compositions typically used in the field (to maximize the chance of comparison to previous studies), modified to reduce RNAse activity and ensure maximal gain of mRNA content.

Electrophysiology signals were recorded using an ITC-18 Data Acquisition Interface (HEKA). Commands were generated, signals processed, and amplifier metadata were acquired using MIES written in Igor Pro (Wavemetrics). Data were filtered (Bessel) at 10 kHz and digitized at 50 kHz. Data were reported uncorrected for the measured (Neher 1992) −14 mV liquid junction potential between the electrode and bath solutions.

Prior to data collection, all surfaces, equipment and materials were thoroughly cleaned in the following manner: a wipe down with DNA away (Thermo Scientific), RNAse Zap (Sigma-Aldrich), and finally nuclease-free water.

After formation of a stable seal and break-in, the resting membrane potential of the neuron was recorded (typically within the first minute). A bias current was injected, either manually or automatically using algorithms within the MIES data acquisition package, for the remainder of the experiment to maintain that initial resting membrane potential. Bias currents remained stable for a minimum of 1 s before each stimulus current injection.

To be included in analysis, a cell needed to have a >1 GΩ seal recorded before break-in and an initial access resistance <20 MΩ and <15% of the R_input_. To stay below this access resistance cut-off, cells with a low input resistance were successfully targeted with larger electrodes. For an individual sweep to be included, the following criteria were applied: 1) the bridge balance was <20 MΩ and <15% of the *R*_input_; 2) bias (leak) current 0 ± 100 pA; and 3) root mean square noise measurements in a short window (1.5 ms, to gauge high frequency noise) and longer window (500 ms, to measure patch instability) were <0.07 mV and 0.5 mV, respectively.

Extracting the nucleus at the conclusion of the electrophysiology experiment led to a substantial increase in transcriptomic data quality. Upon completion of electrophysiological examination, the pipette was centered on the soma or placed near the nucleus (if visible). A small amount of negative pressure was applied (~-30 mbar) to begin cytosol extraction and attract the nucleus to the tip of the pipette. After approximately one minute, the soma had visibly shrunk and/or the nucleus was near the tip of the pipette. While maintaining the negative pressure, the pipette was slowly retracted in the x and z direction. Slow, continuous movement was maintained while monitoring pipette seal. Once the pipette seal reached >1 GΩ and the nucleus was visible on the tip of the pipette, the speed was increased to remove the pipette from the slice. The pipette containing internal solution, cytosol, and nucleus was removed from the pipette holder and contents were expelled into a PCR tube containing lysis buffer (Takara, 634894).

### cDNA amplification and library construction

We used the SMART-Seq v4 Ultra Low Input RNA Kit for Sequencing (Takara, 634894) to reverse transcribe poly(A) RNA and amplify full-length cDNA according to the manufacturer’s instructions. We performed reverse transcription and cDNA amplification for 20 PCR cycles in 0.65 ml tubes, in sets of 88 tubes at a time. At least 1 control 8-strip was used per amplification set, which contained 4 wells without cells and 4 wells with 10 pg control RNA. Control RNA was either Universal Human RNA (UHR) (Takara 636538) or control RNA provided in the SMART-Seq v4 kit. All samples proceeded through Nextera XTDNA Library Preparation (Illumina FC-131-1096) using either Nextera XT Index Kit V2 Sets A-D (FC-131-2001, 2002, 2003, 2004) or custom dual-indexes provided by IDT (IntegratedDNA Technologies). Nextera XT DNA Library prep was performed according to manufacturer’s instructions, except that the volumes of all reagents including cDNA input were decreased to 0.2x by volume. Each sample was sequenced to approximately 500k reads.

### RNA-sequencing

Fifty-base-pair paired-end reads were aligned to GRCm38 (mm10) using a RefSeq annotation gff file retrieved from NCBI on 18 January 2016 (https://www.ncbi.nlm.nih.gov/genome/annotation_euk/all/). Sequence alignment was performed using STAR v2.5.3 (Dobin et al. 2012) in two pass Mode. PCR duplicates were masked and removed using STAR option “bamRemoveDuplicates.” Only uniquely aligned reads were used for gene quantification. Gene counts were computed using the R Genomic Alignments package (Lawrence et al. 2013). Overlaps function using “IntersectionNotEmpty” mode for exonic and intronic regions separately. Exonic and intronic reads were added together to calculate total gene counts; this was done for both the reference dissociated cell data set and the Patch-seq data set of this study.

### Morphological reconstruction

#### Biocytin histology

A horseradish peroxidase (HRP) enzyme reaction using diaminobenzidine (DAB) as the chromogen was used to visualize the filled cells after electrophysiological recording, and 4,6-diamidino-2-phenylindole (DAPI) stain was used identify cortical layers.

#### Imaging

Mounted sections were imaged as described previously (Gouwens et al. 2019). Briefly, operators captured images on an upright AxioImager Z2 microscope (Zeiss, Germany) equipped with an Axiocam 506 monochrome camera and 0.63x optivar. Two-dimensional tiled overview images were captured with a 20X objective lens (Zeiss Plan-NEOFLUAR 20X/0.5) in brightfield transmission and fluorescence channels. Tiled image stacks of individual cells were acquired at higher resolution in the transmission channel only for the purpose of automated and manual reconstruction. Light was transmitted using an oil-immersion condenser (1.4 NA). High-resolution stacks were captured with a 63X objective lens (Zeiss Plan-Apochromat 63x/1.4 Oil or Zeiss LD LCI Plan-Apochromat 63x/1.2 Imm Corr) at an interval of 0.28 μm (1.4 NA objective) or 0.44 μm (1.2 NA objective) along the Z axis. Tiled images were stitched in ZEN software and exported as single-plane TIFF files.

#### Anatomical location

To characterize the position of biocytin-labeled cells in the mouse brain, a 20x brightfield and/or fluorescent image of DAPI-stained tissue was captured and analyzed to determine layer position and region. Soma position of reconstructed neurons was annotated and used to calculate soma depth relative to drawings of the pia and/or white matter. Individual mouse cells were also manually placed in the appropriate cortical region and layer within the Allen Mouse Common Coordinate Framework (CCF) by matching the 20x image of the slice with a “virtual” slice at an appropriate location and orientation within the CCF. Laminar locations were calculated by finding the path connecting pia and white matter that passed through the cell’s coordinate, identifying its distance to pia and white matter as well as position within its layer, then aligning those values to an average set of layer thicknesses. Using the DAPI image, laminar borders were also drawn for all reconstructed neurons.

#### Morphological Reconstruction

Reconstructions were generated based on a 3D image stack that was run through a Vaa3D-based image processing and reconstruction pipeline (Peng et al. 2010). Initial reconstructions were generated with an automated reconstruction of the neuron using TReMAP (Zhou et al. 2015), using reconstruction software PyKNOSSOS (Ariadne-service) or the citizen neuroscience game Mozak (Mozak.science) (Roskams and Popović 2016). Automated or manually-initiated reconstructions were then extensively manually corrected and curated using a range of tools (e.g., virtual finger, polyline) in the Mozak extension (Zoran Popovic, Center for Game Science, University of Washington) of Terafly tools (Bria et al. 2016; Peng et al. 2014) in Vaa3D. Every attempt was made to generate a completely connected neuronal structure while remaining faithful to image data. If axonal processes could not be traced back to the main structure of the neuron, they were left unconnected.

Before morphological feature analysis, reconstructed neuronal morphologies were expanded in the dimension perpendicular to the cut surface to correct for shrinkage (Egger, Nevian, and Bruno 2007; Deitcher et al. 2017) after tissue processing. The amount of shrinkage was calculated by comparing the distance of the soma to the cut surface during recording and after fixation and reconstruction. A tilt angle correction was also performed based on the estimated difference (via CCF registration) between the slicing angle and the direct pia-white matter direction at the cell’s location (Gouwens et al. 2019).

## Data Availability Statement

The data used in this manuscript, the software packages, the detailed protocol, and online resources are freely available to the public and have been consolidated at https://github.com/AllenInstitute/patchseqtools.

## Acknowledgments

We would like to thank the following teams for the services and support. Reagent prep and immunohistochemistry: Medea McGraw, Kris Bickley, Jasmine Bomben, Krissy Brouner, Tom Egdorf, Amanda Gary, Michelle Maxwell, Daniel Park, Alice Pom, and Augustin Ruiz. Human and Mouse Tissue Processing: Nick Dee, Elizabeth Barkan, Tamara Casper, Kristen Crichton, Matt Kroll, Josef Sulc, and Herman Tung. Macaque Tissue Processing: Jonathan Ting, Victoria Omstead, and Natalie Weed. Molecular biology: Darren Bertagnolli, Jeff Goldy, Delissa McMillen, and Michael Tieu. Imaging: Rusty Nicovich, Rachel Enstrom, Melissa Gorham, Maddie Hupp, Samuel Lee, and Lydia Potekhina. Morphology and Reconstructions: Rusty Nicovich, Lauren Alfiler, Alex Henry, Sara Kebede, Matt Mallory, Alice Mukora, David Sandman, Grace Williams and the Mozak citizen-scientists. We are thankful for our collaborator neurosurgeons at the local hospital sites: Charles Cobbs, Richard Ellenbogen, Manuel Ferreira, Ryder Gwinn, Andrew Ko, Jeffrey Ojemann, Akshal Patel, Daniel Silbergeld. We appreciate the constructive feedback on the manuscript provided by Shreejoy Tripathy and Xiao Luo. Funding: NIH grants P51OD010425 (B.E.K.) from the Office of Research Infrastructure Programs (ORIP), UL1TR000423 (B.E.K.) from the National Center for Advancing Translational Sciences (NCATS), and U01 MH114812-02 (E.S.L). This work was funded by the Allen Institute for Brain Science. We wish to thank the Allen Institute founder, Paul G. Allen, for his vision, encouragement, and support.

## Supplementary Material

**Supplementary Table 1.**
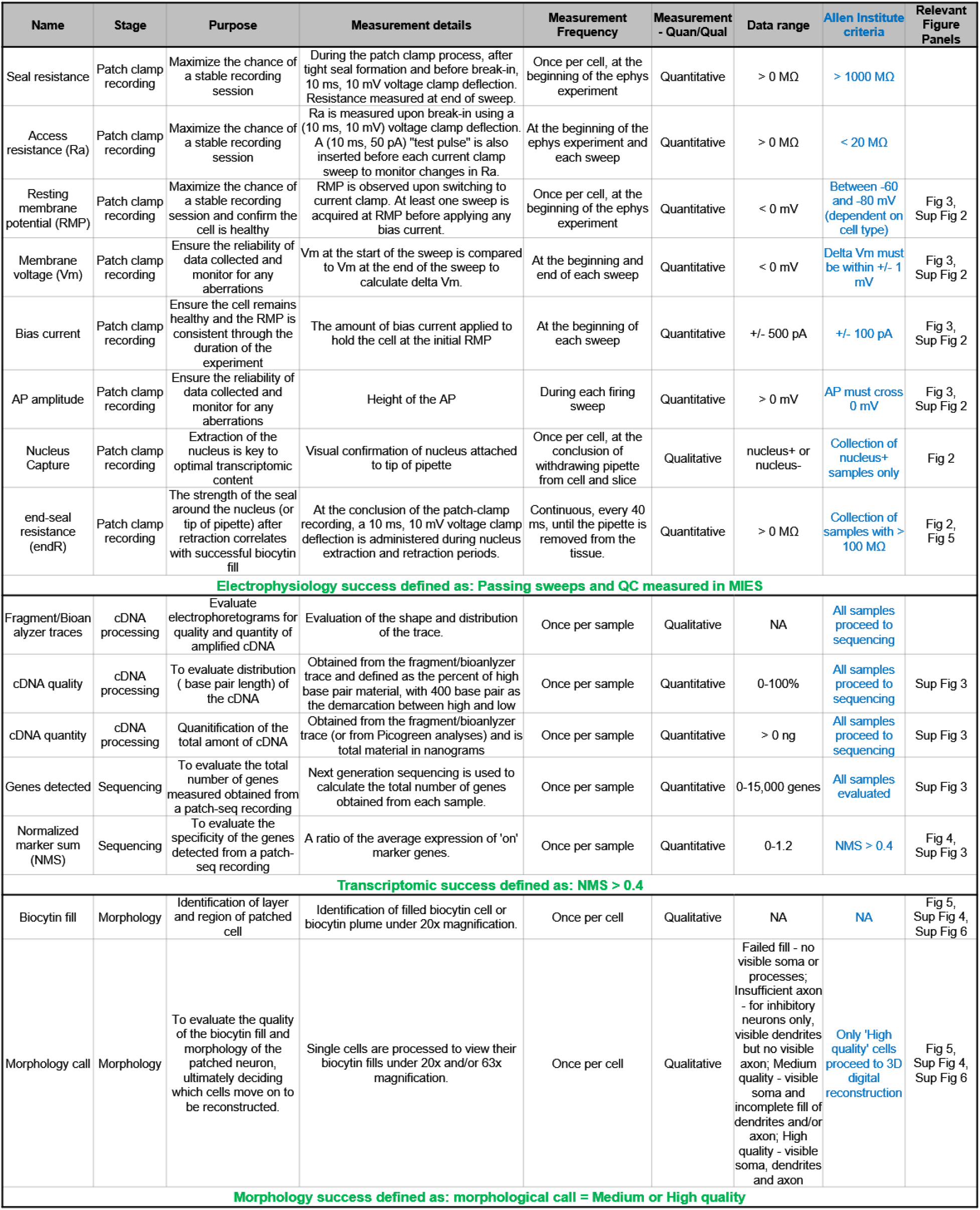
Patch-seq workflow states and QC definitions. The purpose of this table is to list the different parameters and purpose for each stage within the Patch-seq protocols, the metrics measured, the subsequent data and range of data obtained. The criteria that the Allen Institute uses for Patch-seq can be adopted, relaxed, or disregarded.

### Supplementary Figures

**Supplementary Figure 1.**
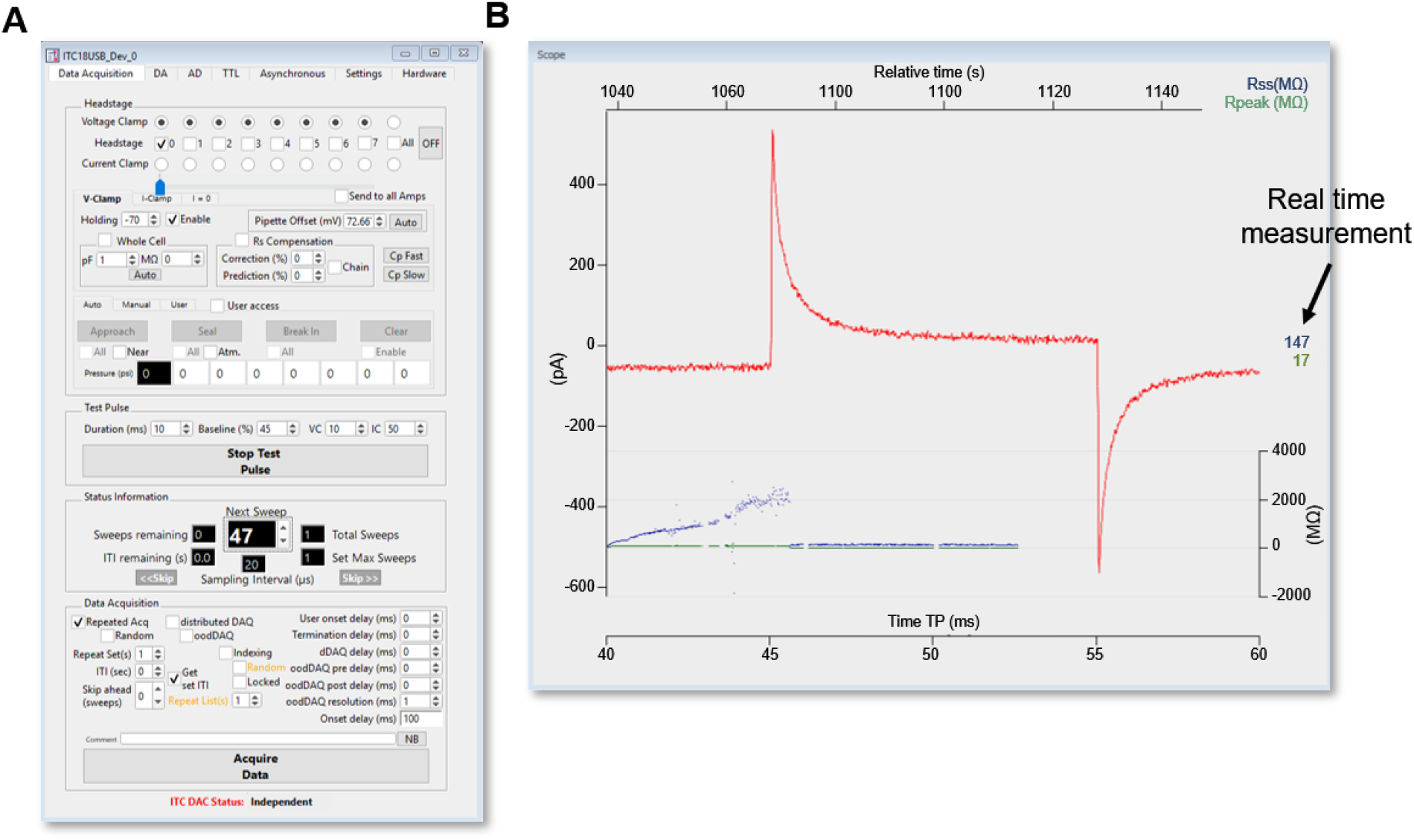
MIES dashboard acquisition. (**A**) MIES Graphical User Interface (GUI) of Data Acquisition. Head stage, Test Pulse, Status Information, Data Acquisition. (**B**) Scope window displays digitized data in real time from test pulse or selected stimulus set(s).

**Supplementary Figure 2.**
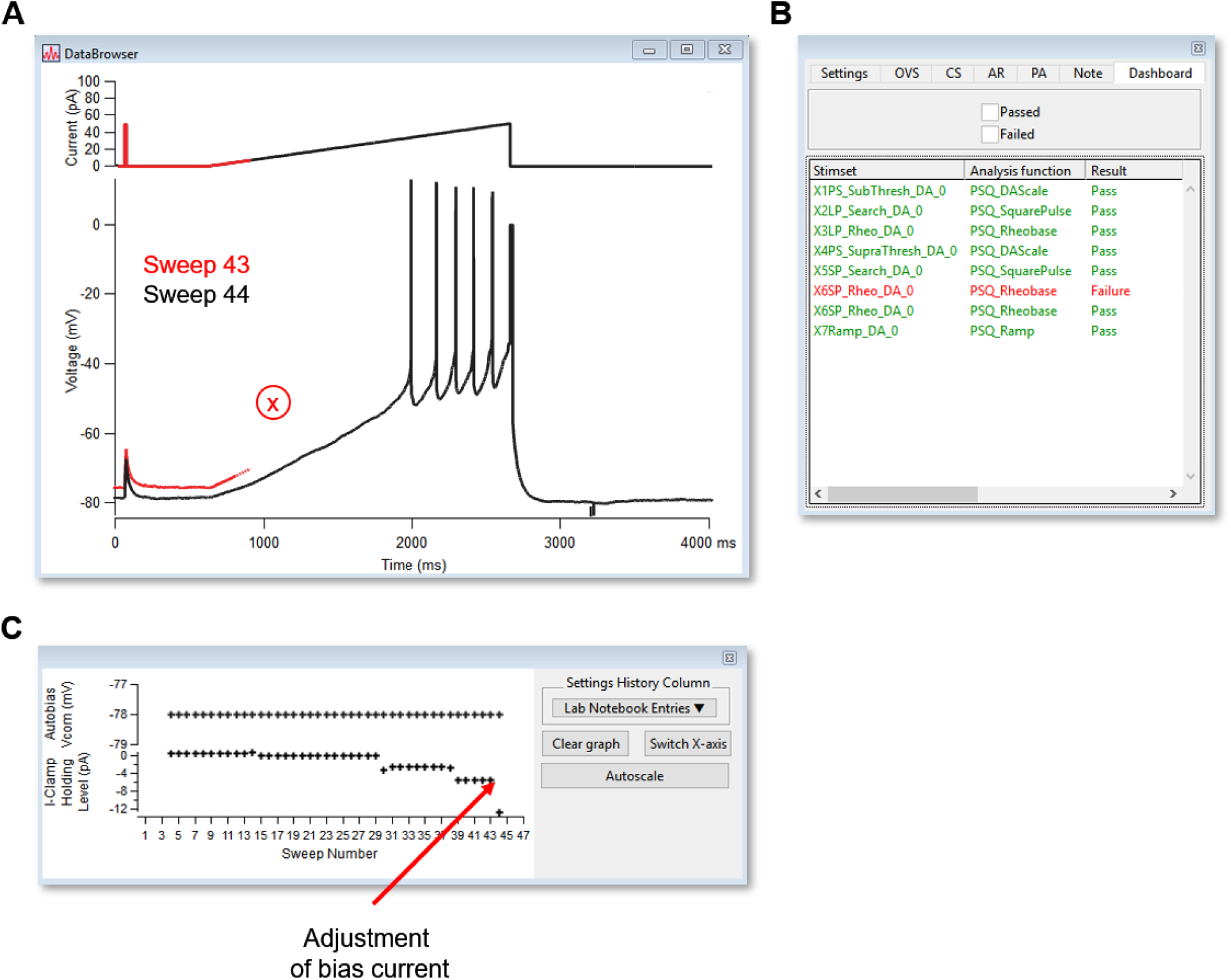
MIES dashboard sweep viewer. (**A**) Data browser/sweep viewer used to display most recent sweep and/or previous sweeps. Example shown is Sweep 43, in red, and its failure to maintain Allen Institute criteria of ±1 mV of designated RMP. Adjustment of Bias current was applied to maintain designated RMP, as noted in (**B**) and sweep 44 was rerun, passing all QC criteria. (**B**) Lab notebook. A variety of features can be plotted during the course of recording. (**C**) Dashboard displaying results of each stimulus set.

**Supplementary Figure 3.**
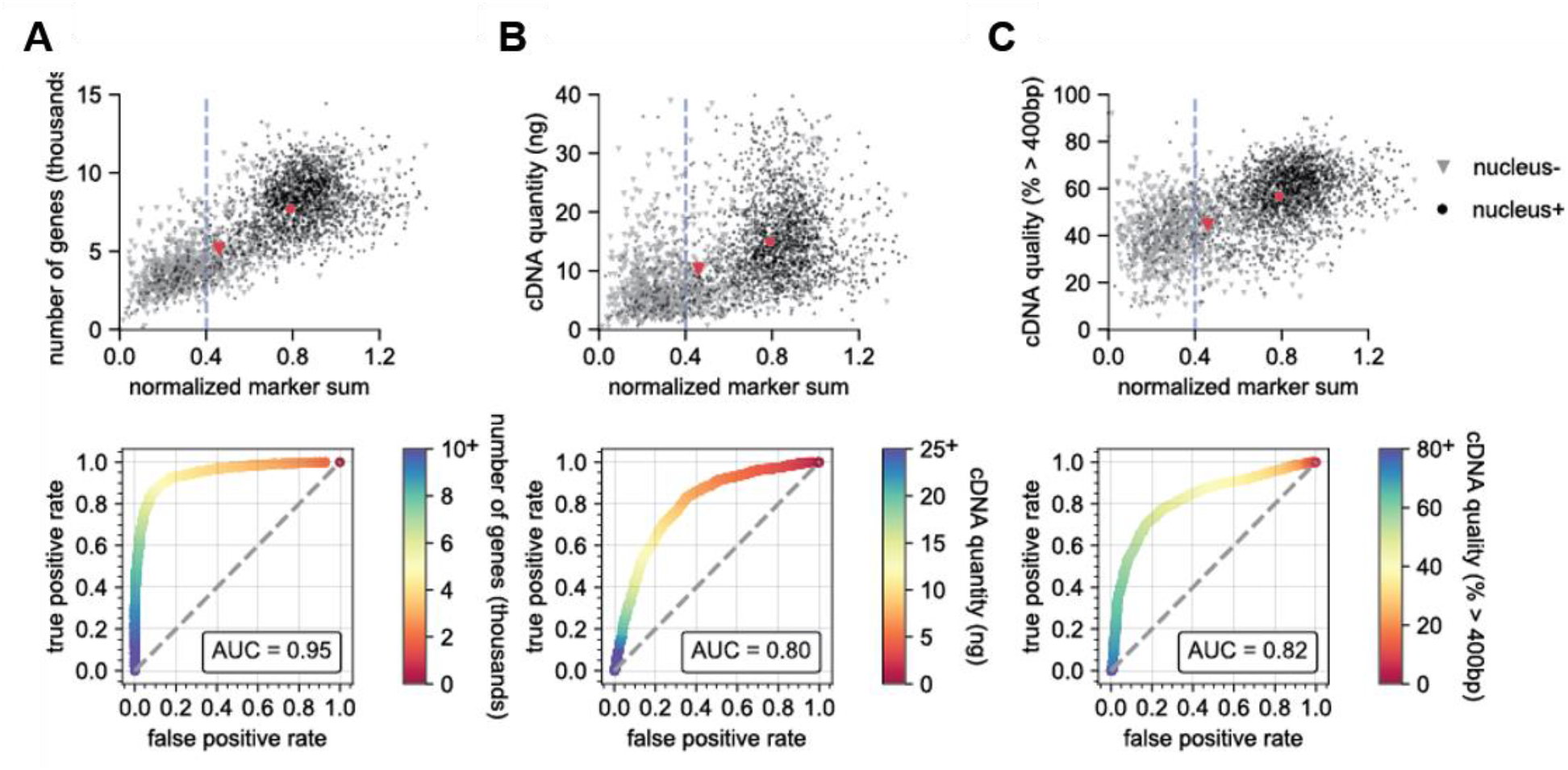
Nucleus+ is predictive for successful sequencing and cDNA data. Scatter plots (top) showing the distribution of values of the NMS and ROC analyses (bottom) of the number of genes detected (**A**), cDNA quantity (ng)(**B**), or cDNA quality (% >400 b.p.) (**C**). In the scatter plots, gray triangles are nucleus- samples, whereas black circles are nucleus+ samples. The larger red symbols represent the means for each group. The dashed blue line represents the NMS pass/fail threshold of 0.4 (Allen Institute criteria). N=1,844 nucleus+; N=952 nucleus- samples

**Supplementary Figure 4.**
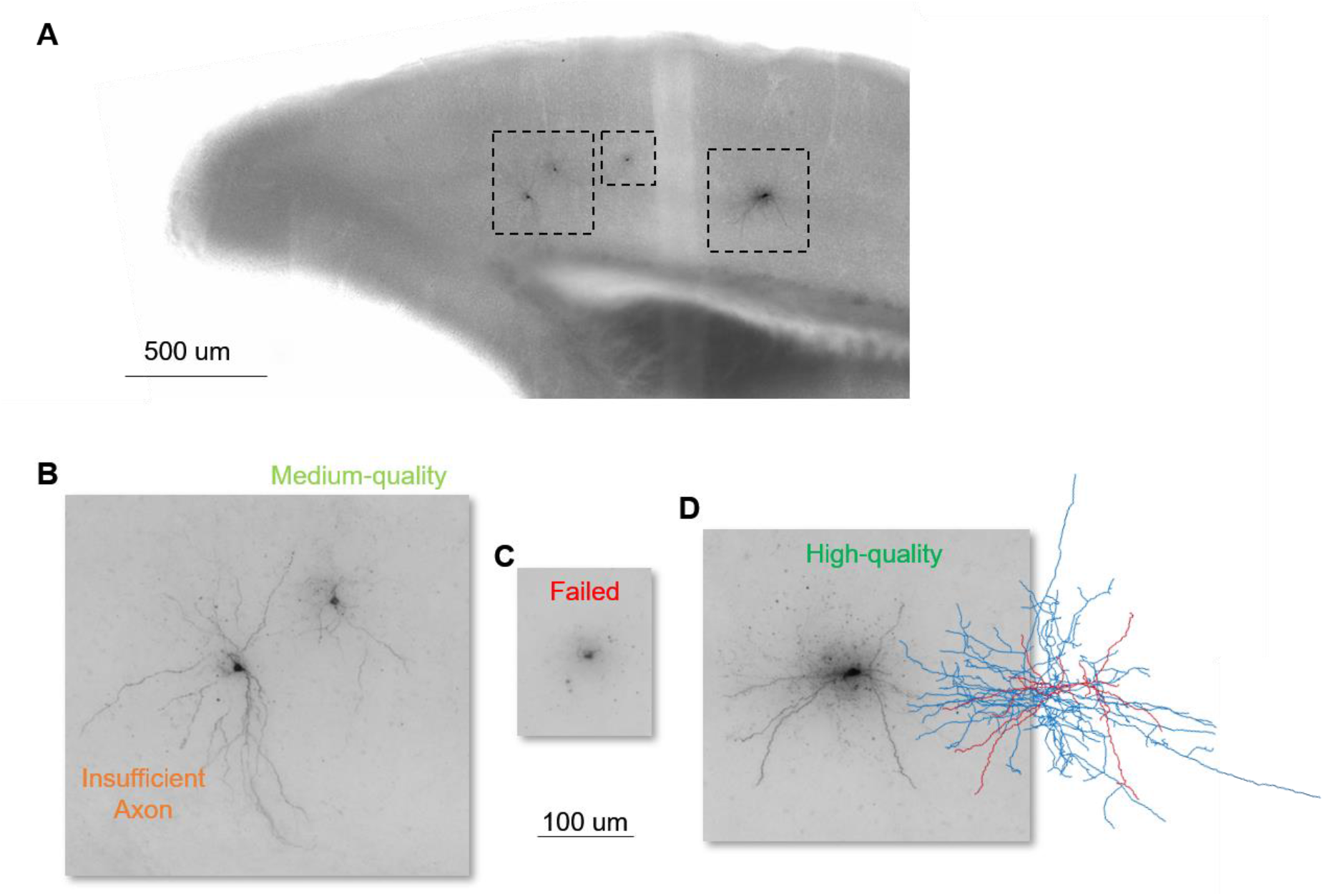
Example biocytin recovery and morphology calls. (**A**) is a 20x brightfield image of a coronal slice from a Sst-Cre line containing biocytin fills from four Patch-seq recordings. (**B-D**) are 63x MIP images from the regions identified in (**A**) and their subsequent morphology outcome. (**B**) contains insufficient axon, left, and medium quality, right, fills; (**C**) is a failed fill, and (**D**) is a high-quality fill with corresponding morphological reconstruction with dendrites in red and axon in blue.

**Supplementary Figure 5.**
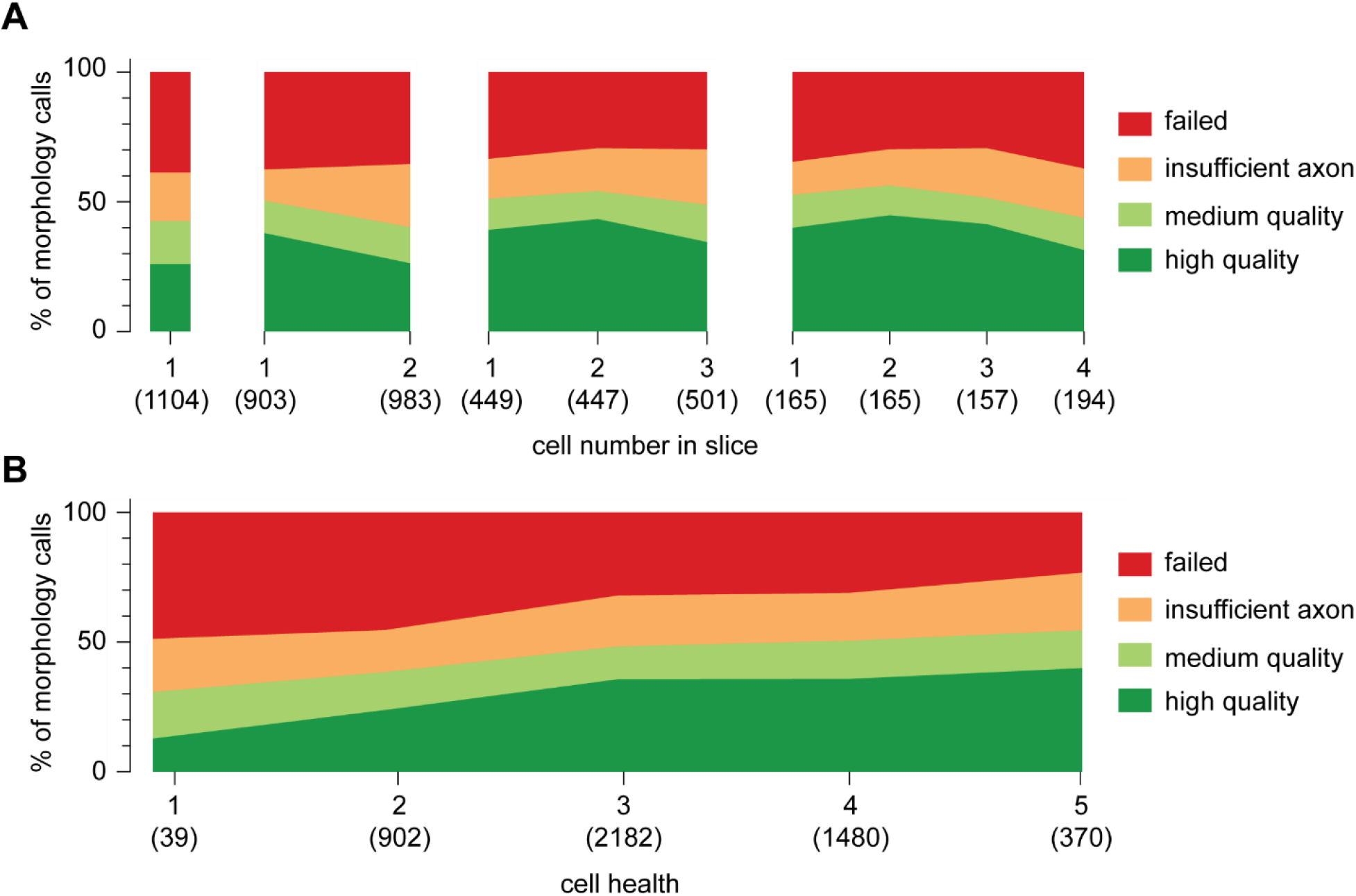
Patching sequence and how it relates to morphological recovery. (**A**) A stacked area plot displaying the percentage of morphology outcomes and how they relate to cell health. Cell health is based on a qualitative assessment of visual appearance and recording quality, with a score of 1 designated as worst ranging to 5 as best. Numbers below, in parentheses, are sample sizes. (**B**) A stacked area plot displaying the percentage of morphology outcomes and how they relate to the cell number in a slice. Blocks shown are slices that had 1, 2, 3, or 4 recordings per slice before fixation in PFA. Numbers below, in parentheses, are sample sizes.

**Supplementary Figure 6.**
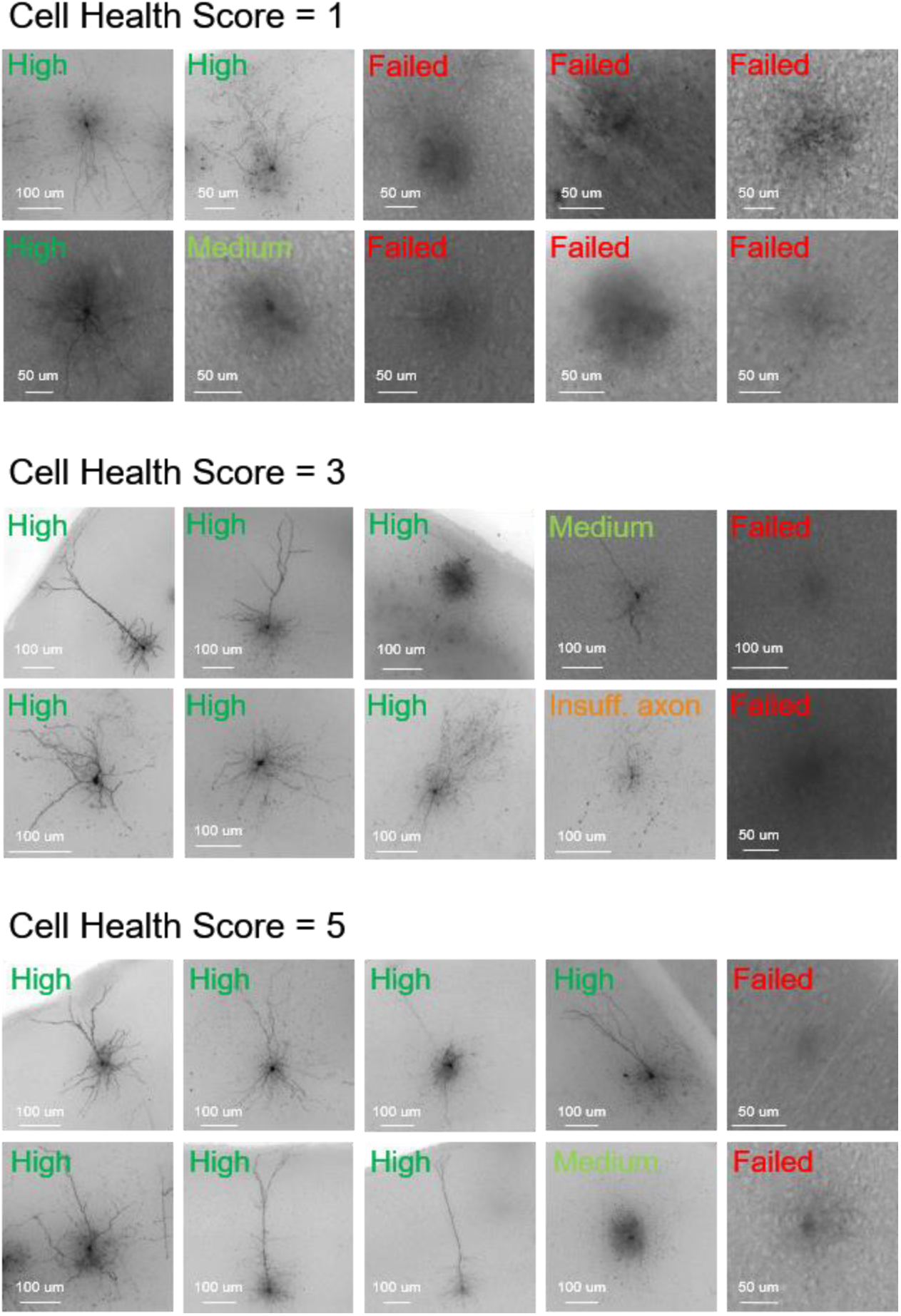
Cell health and morphology outcome examples. Cell health scores of 1, 3, or 5 and high-resolution 63x MIPs of the biocytin-stained fills. Their subsequent morphological outcome is listed in each image. Cell IDs clockwise starting in upper left corner Cell health score 1: 661331024, 750807290, 869512878, 882454517, 652956871, 665602299, 855823375, 816110624, 730878829, 745040951 Cell health score 3: 841854478, 904059133, 914412754, 932264861, 880817724, 898967534, 825552023, 836592858, 875021432, 863585604 Cell health score 5: 761304075, 759948010, 720318238, 823868835, 920542161, 852202575, 809113130, 963287636, 759948010, 831175106

**Supplementary Figure 7.**
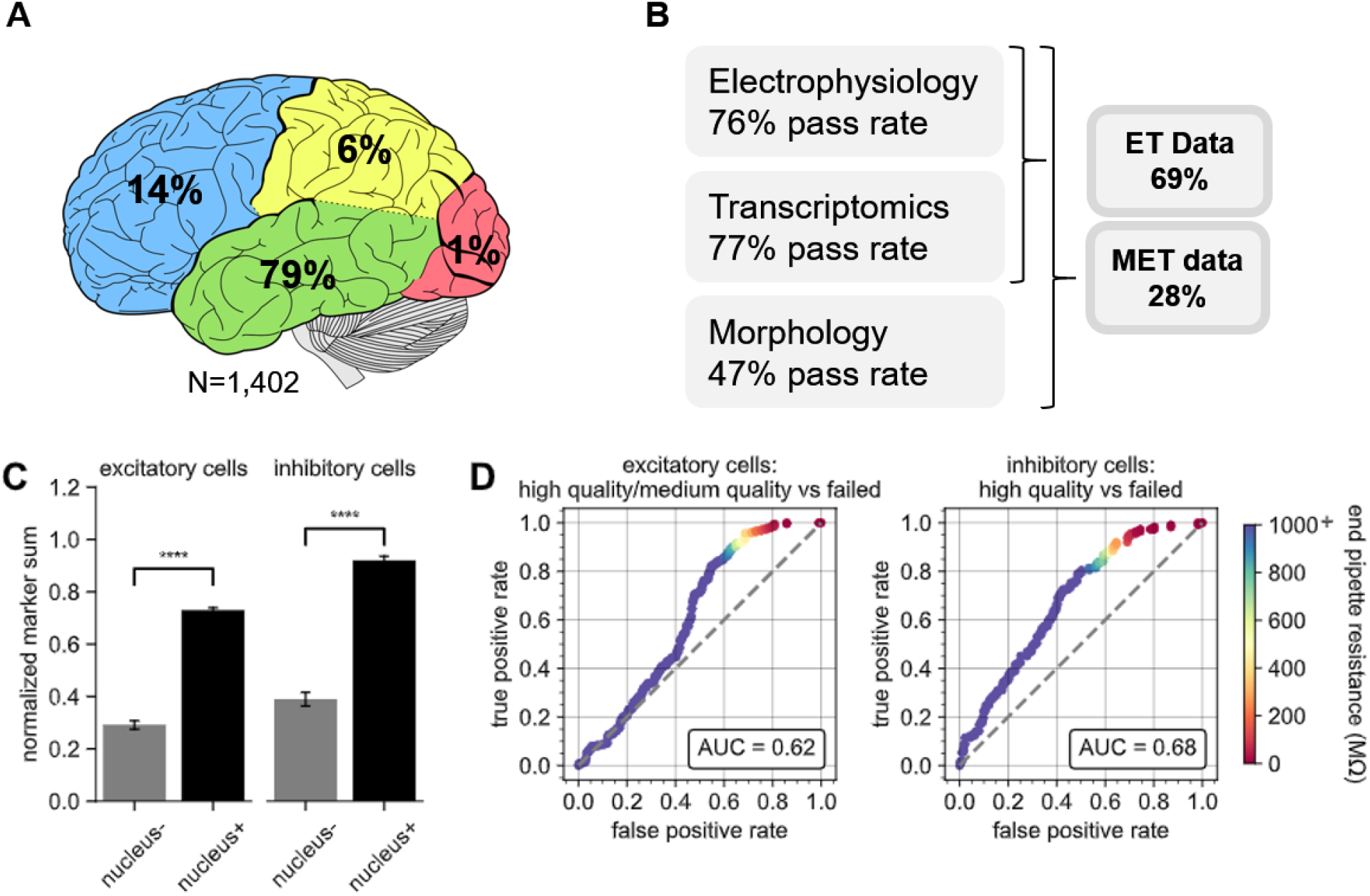
Application of the Patch-seq protocol to human neocortical cells. (**A**) Depiction showing the distribution of human neocortical Patch-seq recordings and the lobes which the tissue was surgically excised from. In (**B**) are the pass rates for each of the independent modalities, ET characterization and MET characterization (N=1,402). The bar plots in (**C**) represent the difference in NMS scores in nucleus- and nucleus+ samples from excitatory N=154 nucleus-; N=668 nucleus+) and inhibitory (N=133 nucleus-; N=462 nucleus+) classes. Nucleus+ have significantly higher NMS scores than nucleus- samples, t(820)=21.23 for excitatory; t(593)=16.38 for inhibitory, p<0.0001 ROC analyses comparing high/medium quality versus failed morphology outcomes for patched (**D**) excitatory (N=212 high/medium quality; N=275 failed) and inhibitory neurons (N=113 high quality; N=223 failed).

**Supplementary Figure 8.**
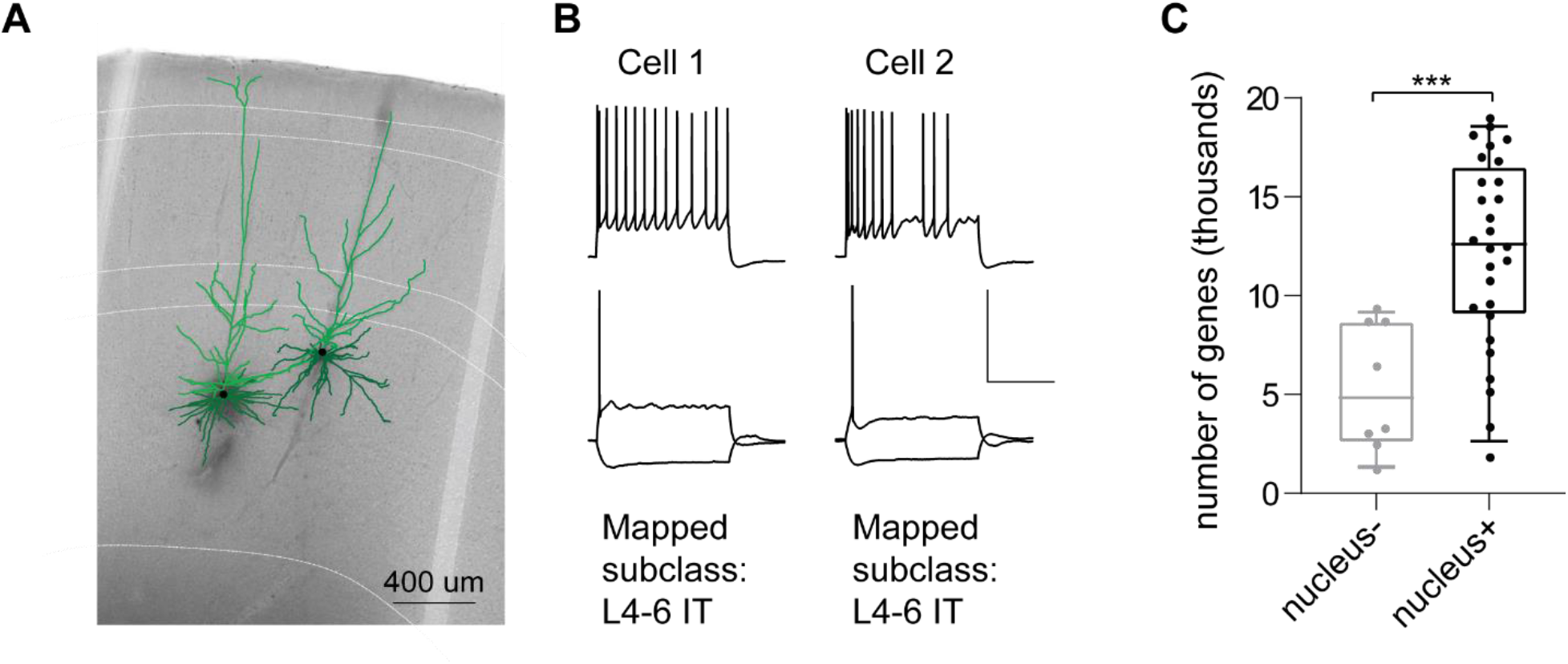
Application of the Patch-seq protocol to macaque neocortical cells. (**A**) is a 20x bright field image of a macaque acute neocortical slice containing biocytin reacted fills with overlaid digital reconstructions from two Patch-seq recordings. Axons are light weight lines and the basal/apical dendrites for excitatory neurons are distinguishable by darker and lighter colors, respectively. Scale bar is 400 um. (**B**) Voltage responses of corresponding cell to 1 s-long current steps equal to −70 pA and rheobase (bottom traces) and rheobase + 80 pA (top trace). Scale bar, vertical 80 mV, horizontal 500 ms. Both cell 1 and 02 mapped to the excitatory subclass L4-6 IT. (**C**) Box plots showing the distribution of values for the number of genes detected for nucleus- and nucleus+ samples, t(34)=3.80, p<0.001.

## Notes

### Competing Interest Statement

The authors have declared no competing interest.

